# Evolution of the Analytical Scattering Model of Live *Escherichia Coli*

**DOI:** 10.1101/2020.09.18.303305

**Authors:** Enrico F. Semeraro, Lisa Marx, Johannes Mandl, Moritz P. K. Frewein, Haden L. Scott, Sylvain Prévost, Helmut Bergler, Karl Lohner, Georg Pabst

## Abstract

We have revised a previously reported multi-scale model for (ultra) small angle X-ray (USAXS/SAXS) and (very) small angle neutron scattering (VSANS/SANS) of live *Escherichia coli* based on compositional/metabolomic and ultrastructural constraints. The cellular body is modelled, as previously described, by an ellipsoid with multiple shells. However, scattering originating from flagella was substituted by a term accounting for the oligosaccharide cores of the lipopolysaccharide leaflet of the outer membrane including its cross-term with the cellular body. This was mainly motivated by (U)SAXS experiments showing indistinguishable scattering for bacteria in the presence and absence of flagella or fimbrae. The revised model succeeded in fitting USAXS/SAXS and differently contrasted VSANS/SANS data of *E. coli* ATCC 25922 over four orders of magnitude in length scale, providing specifically detailed insight into structural features of the cellular envelope, including the distance of the inner and outer membranes, as well as the scattering length densities of all bacterial compartments. Consecutively, the model was also successfully applied to *E. coli* K12, used for our original modelling, as well as for two other *E. coli* strains, detecting significant differences between the different strains in terms of bacterial size, intermembrane distance and its positional fluctuations. These findings corroborate the general applicability of our approach to quantitatively study the effect of bactericidal compounds on ultrastructural features of Gram-negative bacteria without the need to resort to any invasive staining or labelling agents.

## 1. Introduction

*E. coli* are among the most studied Gram-negative bacterial strains in life-sciences, with numerous reports on its structure and composition (Breed & Dotterrer, 1916; Lieb *et al*., 1955; Maclean & Munson, 1961; Neidhardt *et al*., 1990; Seltmann & Holst, 2002; Silhavy *et al*., 2010). Transmission electron microscopy (TEM) has been an indispensable tool to derive the cell’s ultrastructure, i.e. the few tens of nanometer thick structure of the bacterial cell wall (Milne & Subramaniam, 2009). It took, however, significant efforts to minimize limitations originating from the invasive nature of the technique (Hobot *et al*., 1984; Matias *et al*., 2003). Nevertheless, TEM still evades dynamic studies of the cellular ultrastructure under physiological relevant conditions and thus real-time insight on its modification by bactericidal compounds, such as antimicrobial peptides. Time-resolved small-angle scattering experiments (Huxley *et al*., 1980), capable of probing structural heterogeneities on the (sub)micrometer to subnanometer length scales without the need of using either bulky labels or invasive staining techniques, are possible solutions for this issue (see, e.g. (Zemb & Lindner, 2002) for an updated overview on scattering techniques). In terms of static, equilibrium experiments, small-angle neutron scattering (SANS) has been used, for example, to probe the response of thylakoid membranes in live chloroplasts to external stimuli (Liberton *et al*., 2013; Nagy *et al*., 2014), while a principle component analysis of small-angle X-ray scattering (SAXS) data was applied to get some qualitative insight on the effect of antibiotics on *E. coli* (von Gundlach *et al*., 2015; von Gundlach *et al*., 2016).

Quantitative insight on live cell’s ultrastructure using either SAXS or SANS is challenging, however. This is simply due to the fact that both techniques provide a global average of entire cellular content, making attempts to single-out individual contributors a veritable tour de force (Semeraro *et al*., 2020). This can be addressed, upon extensive use of the contrast variation capabilities of SANS. For example Nickels et al. (Nickels *et al*., 2017) were able to grow fully deuterated *Bacillus subtilis* fed with mixtures of protiated and deuterated fatty acids, which allowed them to highlight nanoscopic domains within the bacteria’s cytoplasmic membrane using SANS. Yet, it is also possible to obtain insight on cellular ultrastructure without the need of growing bacteria in D_2_O. In particular, we recently reported a multi-scale model that successfully describes the scattered intensities of live *E. coli* K12 originating from ultra-SAXS (USAXS), SAXS and SANS experiments at five D_2_O/H_2_O ratios (Semeraro *et al*., 2017). Specifically, the model applies a core-shell description of the cell’s body, composed of an ellipsoidal cytoplasmic space, a multilayered cellular wall, and includes contributions from flagella in terms of self-avoiding polymer chains (Doi & Edwards, 1988). The latter term was found to significantly contribute at intermediate to high magnitudes of the scattering vector **q**.

Continuing our efforts to use elastic scattering techniques for exploiting the ultrastructure of *E. coli* led us to perform SAXS experiments on *E. coli* ATCC 25922 with either regular or short flagella, and the ATCC flagella-free mutant Δ*fliC* with the surprising result of basically superimposable scattering patterns (see Fig. S1, in the supporting information). This prompted us to thoroughly revise our analytical form factor model based on robust estimates of the molecular composition of the bacteria and their structural integration. These estimates include sizes, volumes, concentrations and distribution of all major bacterial components.

Briefly, the most important changes of our revised bacterial model are: *i*) constraints for the average scattering length densities (SLDs) of different cell compartments were derived by considering their constituting macromolecules as separate bodies, including estimates of SLDs of the metabolome and membrane; *ii*) contributions from flagella were substituted by the oligosaccharide cores of lipopolysaccharide (LPS), modelled as grafted polymers; and *iii*) variations of the inter-membrane distance are modelled by a log-normal probability distribution function (PDF) along with removing the negligible polydispersity over the cell radius from the model.

The model was tested against USAXS/SAXS and Very-SANS (VSANS)/SANS data at ten different contrasts of *E. coli* ATCC 25922 yielding highly satisfactory fits over the complete range of recorded scattering vector magnitudes (3 *×* 10^*−*3^ nm^*−*1^ *< q <* 7 nm^*−*1^). These tests include also a more complex model, considering a heterogeneously structured cytosol and including specifically the scattering originating from ribosomes. Our analysis showed, however, that its contribution to overall scattering is overwhelmed by that of the cell wall. The new model was also successfully applied to USAXS/SAXS data of *E. coli* K12, previously used for devising our multi-scale model (Semeraro *et al*., 2017), as well as the fimbrae-free JW4283 and the strain Nissle 1917, reporting distinct differences in ultrastructural features. The paper is structured as follows. First we briefly summarize the experimental methods and samples, before we detail the revised modelling in section 3, including a comprehensive list of compositional data in the supporting information. Section 4 describes an analytical model for scattering from a heterogeneous cytosol accounting for ribosomes and section 5 summarizes the involved parameters and applied optimization strategy. Results of applying the modelling to experimental data of five different *E. coli* strains are described and discussed in section 6, before we conclude in section 7.

## 2. Materials and methods

### 2.1. Bacterial samples

Bacterial colonies of *Escherichia coli* strains ATCC 25922, K12 5K, K12 JW4283 (Baba *et al*., 2006) and Nissle 1917 (Sonnenborn, 2016), were grown in LB-agar (Roth) plates at 37 *°*C. Overnight cultures (ONCs) were derived from these colonies by inoculating a single colony in 3 ml LB-medium (Luria/Miller, Roth) in sterile polypropylene conical tubes (15 ml), allowing for growth under aerobic conditions for 12 *−* 16 hours in a shaking incubator at 37 *°*C. Main cultures (MCs) were then prepared by suspending an aliquot of the ONCs in 10 ml LB-medium in 50 ml sterile polypropylene conical tubes, allowing for bacterial growth under the same conditions applied to ONCs up to the middle of the exponential growth phase. Cells were then immediately washed twice and re-suspended in nutrient-free and isotonic phosphate-buffered saline (PBS) solution (phosphate buffer 20 mM, NaCl 130 mM) at pH 7.4 (Sigma Aldrich). Turbidity measurements were used to control the bacterial concentration. Optical density values at wavelength *λ* = 600 nm (OD_600_) were acquired with the spectrophotometer Thermo Spectronic Genesys 20 (OD_600_ = 1 *≃* 8 *×* 10^8^ colony-forming-units (CFU) per milliliter). In these samples, 1 CFU corresponds to one single cell. In the case of samples containing D_2_O, bacterial suspensions were washed twice with either PBS or 90 wt% D_2_O PBS solutions, in order to obtain two concentrated stock solutions for both buffer conditions. These two stocks were then mixed and diluted down to the required bacterial concentrations and D_2_O contents according to the experimental settings.

The preparation of the ATCC samples was conducted with the maximum care in order to preserve the integrity of the flagella. ATCC cells with mechanically fragmented flagella were prepared by shearing the suspension 5 times through a 22-gauge needle equipped on 3 ml syringe, as described in (Turner *et al*., 2012). The suspension was then washed in PBS to eliminate flagella fragments in the supernatant.

The ATCC 25922 Δ*fliC* strain was constructed by phage transduction as described in (Silhavy *et al*., 1984). The P1vir phage was propagated on strain YK4516 (Komeda *et al*., 1980) using the plate lysate method. YK4516 contains a Tn10 insertion in *fliC* (fliC5303::Tn10) and was purchased from The Coli Genetic Stock Center (CGSC), Yale. Strain ATCC 25922 served as recipient. Transductands were selected on tetracycline containing plates (10 *µ*g/ml) and the *fli*^*−*^ phenotype was confirmed by spotting on semisolid agarplates (0.3% agarose).

### 2.2. Experimental set-up

#### USAXS

USAXS/SAXS measurements were performed on the TRUSAXS beamline (ID02) at the ESRF, Grenoble, France. The instrument uses a monochromatic beam that is collimated in a pinhole configuration. Measurements were performed with a wavelength of 0.0995 nm, and sample-to-detector distances of 30.8, 10.0 and 1.0 m, covering a *q*-range of 0.001 *−* 7 nm^*−*1^ (Narayanan *et al*., 2018). The measured two-dimensional scattering patterns were acquired on a Rayonix MX170 detector, normalized to absolute scale and azimuthally averaged to obtain the corresponding one-dimensional USAXS/SAXS profiles. The normalized cumulative background from the buffer, sample cell and instrument were subtracted to obtain the final *I*(*q*). Samples with bacterial concentration *∼* 10^10^ cells/ml were measured at 37 *°*C and contained in a quartz capillaries of 2 mm diameter, mounted on a flow-through set-up in order to maximize the precision of the background subtraction.

#### VSANS

VSANS/SANS measurements were acquired on D11 instrument at the Institut Laue-Langevin, Grenoble, France, with a multiwire ^3^He detector of 256 *×* 256 pixels (3.75 *×* 3.75 mm^2^). Four different set-ups (sample-to-detector distances of 2, 8, 20.5, and 39 m with corresponding collimations of 5.5, 8, 20.5 and 40.5 m, respectively), at a wavelength *λ* = 0.56 nm (Δ*λ/λ* = 9%), covered a *q*-range of 0.014 *−* 3 nm^*−*1^. To reach very low *q*, a combination of large wavelength (*λ* = 2.1 nm), two focusing MgF_2_ lenses and a mirror to cancel deleterious gravity effects (loss of neutrons in the collimation and loss of resolution due to gravity smearing on the detector) were used; the setup is described in (Cubitt *et al*., 2011). Samples (concentration *∼* 10^10^ cells/ml) were measured at 37 *°*C and contained in quartz Hellma 120-QS banjo-shaped cuvettes of 2 mm pathway. They were mounted on a rotating sample holder, which prevented the bacteria from sedimenting. Data were reduced with the Lamp program from ILL, performing flat field, solid angle, dead time and transmission correction, normalizing by incident flux (via a monitor), and subtracting the contribution from an empty cell. Experimental set-up and data are available in https://doi.ill.fr/10.5291/ILL-DATA.8-03-910.

#### In-house SAXS

A SAXSpace compact camera (Anton Paar, Graz, Austria) equipped with an Eiger R 1 M detector system (Dectris, Baden-Daettwil, Switzerland) was used for laboratory SAXS experiments. Cu-K*α* (*λ* = 1.54 Å) X-rays were provided by a 30 W-Genix 3D microfocus X-ray generator (Xenocs, Sassenage, France). Samples were taken-up in glass capillaries (diameter: 1 mm; Anton Paar) and equilibrated at 37 *°*C for 10 minutes prior to measurement using a Peltier controlled sample stage (TC 150, Anton Paar). The total exposure time was 30 minutes (6 frames of 5 min), setting the sample-to-detector distance to 308 mm. Data reduction, including sectorial data integration and corrections for sample transmission and background scattering, was performed using the program SAXSanalysis (Anton Paar).

#### Dynamic light scattering

Measurements were performed using a Zetasizer Nano ZSP (Malvern Panalytical, UK). ATCC 25922 and K12 5K samples (concentration *∼* 10^7^ cells/ml) were suspended in glass cuvettes of 1 cm path-length and equilibrated at 37 *°*C. This bacterial concentration provided an optimal noise-to-signal ratio and avoided multiple-scattering effects. Data were automatically analyzed by the supplied software (Malvern), which, via a standard cumulant analysis, provided a monomodal probability distribution of the bacterial population as a function of the hydrodynamic radius *R*_*H*_. Each distribution had an average polydispersity index of *∼* 0.25, due to the variations of the cell lengths during growth and division. Average *R*_*H*_ values and the associated errors were calculated from 18 measurements (6 frames of 3 different sample volumes). The absence of energy sources in PBS and vigorous vortexing of the samples (fragmentation of flagella) minimized cell motility, making Brownian random-walk the dominant dynamics of this sample.

## 3. Overall scattering contributions and compositional modelling

A holistic description of elastic scattering from complex Gram-negative prokaryotic cells can be derived by considering first their prevalent molecular and supramolecular components, each of them having a well defined range of lengths, volumes and densities, which then serve as a guide to construct a comprehensive scattering form-factor model. The here applied scattering contribution estimates are based on the latest experimental and computational reports on *E. coli*, including isolated cell components. Specifically, we used compositional and structural information about *E. coli* and its cell-wall (Neidhardt *et al*., 1990; Seltmann & Holst, 2002; Schwarz-Linek *et al*., 2015); the cytoplasmic space and its components (Zimmerman & Trach, 1991; Tweeddale *et al*., 1998; Maharjan & Ferenci, 2003; Bennett *et al*., 2009; Guo *et al*., 2012; Lebedev *et al*., 2015); the bacterial ultrastructure (Hobot *et al*., 1984; Beveridge, 1999; Matias *et al*., 2003); the lipid membrane composition and structure (De Siervo, 1969; Oursel *et al*., 2007; Lohner *et al*., 2008; Pandit & Klauda, 2012; Kučerka *et al*., 2012; Kučerka *et al*., 2015; Leber *et al*., 2018); the LPS specifics (Heinrichs *et al*., 1998; Müller-Loennies *et al*., 2003; Kučerka *et al*., 2008; Kim *et al*., 2016; Rodriguez-Loureiro *et al*., 2018; Micciulla *et al*., 2019); the periplasmic space (Burge *et al*., 1977; Labischinski *et al*., 1991; Pink *et al*., 2000; Gan *et al*., 2008); and the external bacterial components (Yamashita *et al*., 1998; Whitfield & Roberts, 1999; Stukalov *et al*., 2008; Turner *et al*., 2012). All this information has been condensed into SLDs *ρ*, which are summarized in Fig. 1. For a detailed description, please see the supporting information (SI).

**Fig. 1.**
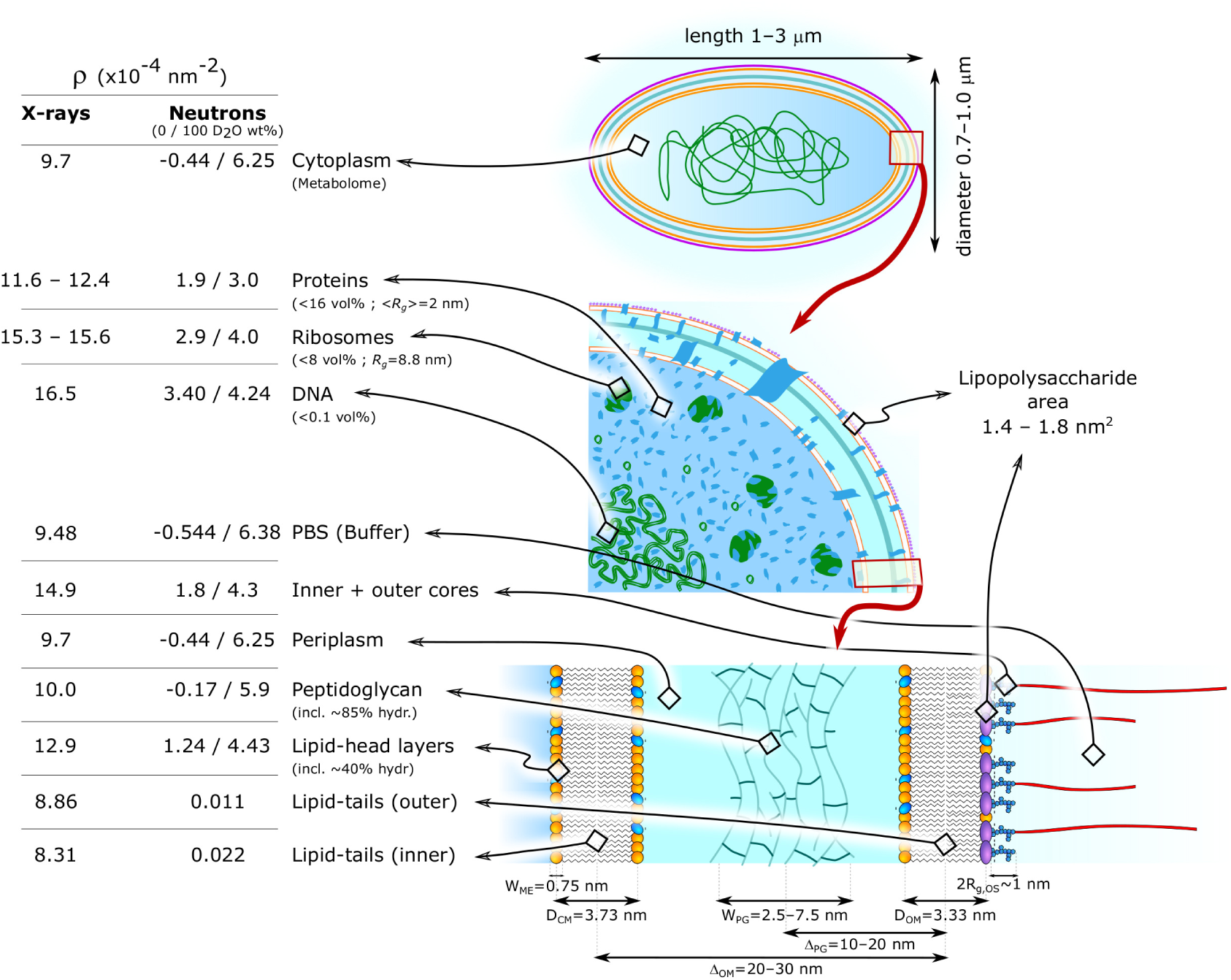
Schematic of *E. coli* structure and composition, including typical sizes, as well as X-ray and neutron SLDs of the most relevant constituents. The bacterial shape is conveniently modeled by an ellipsoid (Semeraro *et al*., 2017).

The benefit of this detailed description becomes clear considering the ability of SANS to nullify or enhance contrast for a given molecular entity, depending on the applied H_2_O/D_2_O ratio. For homogeneous scatterers in a dilute regime, i.e. when their volume fraction *φ ≪* 1, the forward scattering intensity, *I*(0), is related to the scattering invariant, *Q*, as the following:

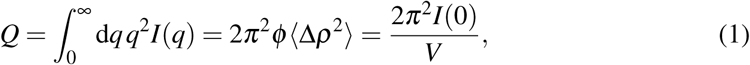

where *V* is the volume of the scatterer, and *⟨*Δ*ρ⟩* is the average scattering contrast (Porod, 1982). For inhomogeneous systems, such as complex live cells, the average contrast can be calculated as a volume-fraction-weighted average of the contrast of each cell compartment/species 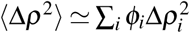, where *φ*_*i*_ is the volume-fraction of the i’th component. Hence, the estimates *φ*_*i*_ and Δ*ρ*_*i*_ enable us to approximate *Q* for all bacterial components (Fig. 2), see also (Nickels *et al*., 2017). This strong approximation leads to a “matching” point of the entire cell at about 40 wt% D_2_O. At higher D_2_O content the total scattering intensity is increasingly dominated by the acyl-chains of the membrane lipids, because they are devoid of water. Toward lower D_2_O contents, cytoplasmic components, such as ribosomes and proteins, are the dominating scattering contributors in turn.

**Fig. 2.**
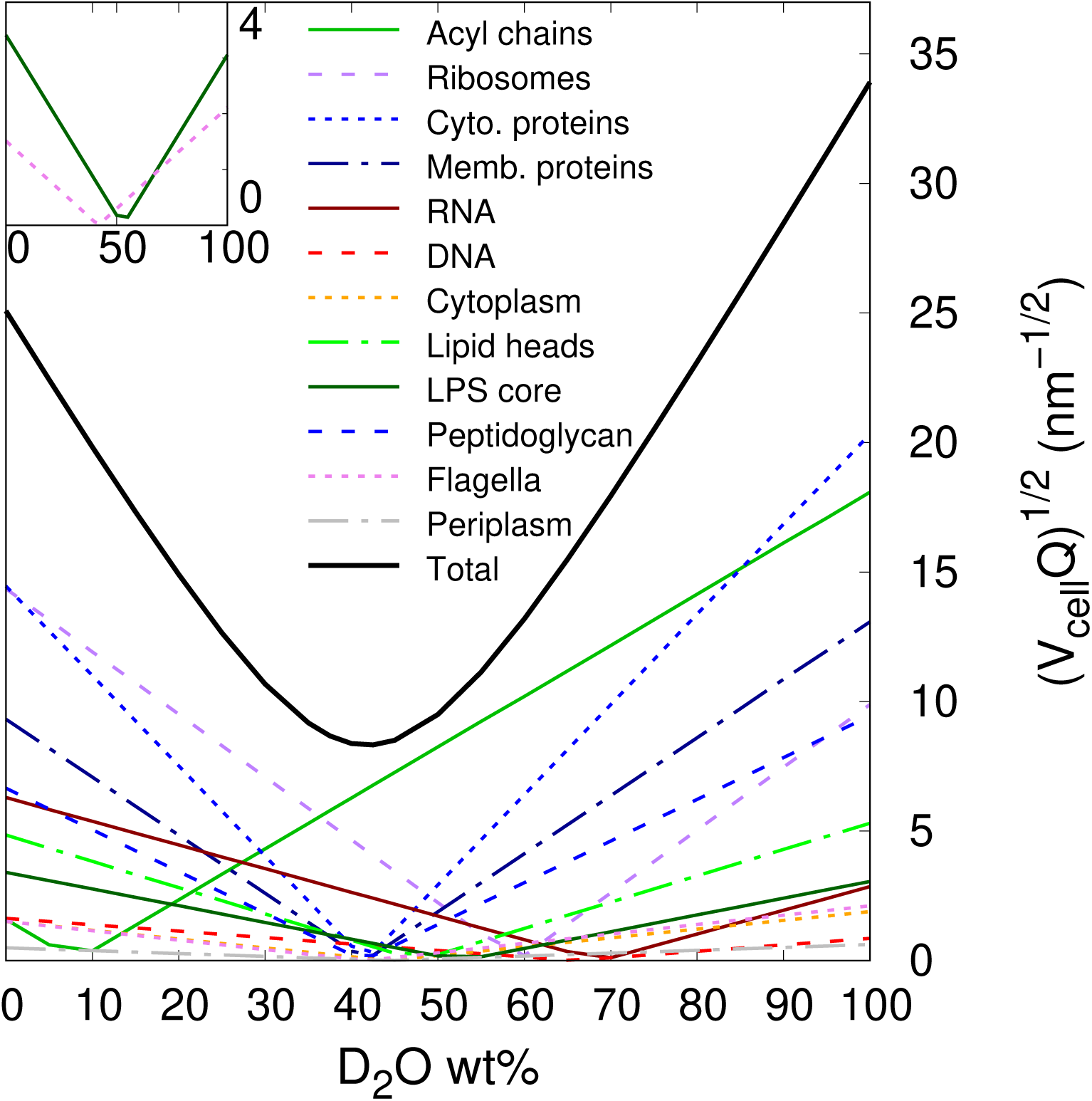
Square root of the estimated Porod invariant *Q* as a function of D_2_O wt%, calculated for component using Eq. (1) and multiplied by the cell volume. The inset marks the differences between the contribution of the LPS oligosaccharide cores (solid green) and flagella (dashed pink).

### 3.1. Multi-scale scattering model

As reported previously (Semeraro *et al*., 2017), the main body of *E. coli* is described in terms of a scattering amplitude of an ellipsoid with multiple shells (multi-core-shell):

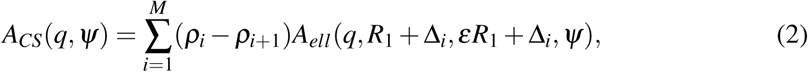

where the *ρ*_*i*_’s are the SLDs given by the compositional *E. coli* averages of each shell of width Δ_*i*_ (*ρ*_*M*+1_ is the SLD of the buffer); *R*_1_ is the radius of the cytoplasmic core; *Ψ* is the angle related to every possible orientation of a prolate in suspension and *ε >* 1 is the ratio of the minor, *R*, and major radii, *εR*, of the ellipsoid. Further,

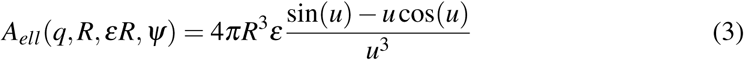

is the scattering amplitude of an ellpsoid (Pedersen, 1997) and

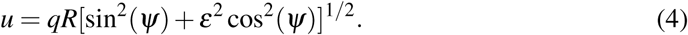

A significant difference to the previous model (Semeraro *et al*., 2017) relates to the *ρ*_*i*_’s used as fitting constrains. Here, we considered, given the available *q*-range, scattering contributions from every macromolecular species (proteins, ribosomes, DNA, etc.) individually, for calculating the average SLDs. This affects, compared to our previously used values, in essence the *ρ*_*i*_ estimates of the cytoplasm – now based on metabolomic analysis – and the phospholipid membranes (Fig. 1). Of note and compared to elastic scattering experiments on lipid-only mimics, structural parameters of each single bilayer cannot be resolved in the context of a whole cell analysis (Semeraro *et al*., 2020). Hence, the Δ_*i*_ and *ρ*_*i*_ values of both membranes were considered as fixed parameters (see Fig. 1, more details about Δ_*i*_ and *ρ*_*i*_ are described in the SI).

Note that the volume fraction of the bacterial suspensions was *≤* 0.007, hence the presence of an inter-cellular structure factor of interactions is unlikely.

Arguably the most distinct differences compared to the former model result from flagella, whose scattering was previously added in terms of a polymer-like structure factor yielding significant contributions to *I*(*q*) for *q >* 0.06 nm^*−*1^ (Semeraro *et al*., 2017). Surprisingly, however, a comparison of SAXS data of native ATCC 25922, ATCC with physically broken, short flagella (Turner *et al*., 2012), as well as the flagella-free Δ *f liC* ATCC mutant, showed indistinguishable scattering intensities in the *q*-range previously thought to be dominated by flagella (Fig. S1). Consequently, flagella do not contribute significantly to the *E. coli* scattering signal. Note that the integrity of flagellar filaments in common bacterial cultures is not guaranteed, as excessive centrifuging or careless sample manipulation steps easily lead to their fragmentation (Schwarz-Linek *et al*., 2015). Even if we cannot guarantee that the reference ATCC sample possessed fully intact flagella, the bacterial suspensions used in this work were prepared with utmost care. Note that in our hands the same sample preparation protocol allowed us to obtain motile bacteria (Semeraro *et al*., 2018), suggesting that the flagellar integrity was preserved at least to some degree.

Attempting to rectify the missing scattering intensities in our multi-scale model led us to consider contributions originating from the oligosaccharide (OS) inner and outer cores. Initially, the function *A*_*CS*_ was modified by a new shell describing the OS-cores, which resulted in nonphysical results, however (Fig. S2). In particular, *ρ*_X-ray_ *∼* 15 *×* 10^*−*4^ nm^*−*2^ of the OS-layer suggested that water is expelled, which is inconsistent with neutron reflectometry experiments on supported LPS layers showing that the hydration of the inner and outer cores ranges from 40 to 80 vol% (Clifton *et al*., 2013; Rodriguez-Loureiro *et al*., 2018; Micciulla *et al*., 2019). We therefore decided to model the OS-cores in terms of grafted blocks on the outer cell surface. Each core was approximated by a Gaussian chain polymer, entailing the application of the polymer-grafted colloid formalism (Pedersen & Gerstenberg, 1996). The scattering form factor of such a system is given by (Pedersen, 2000):

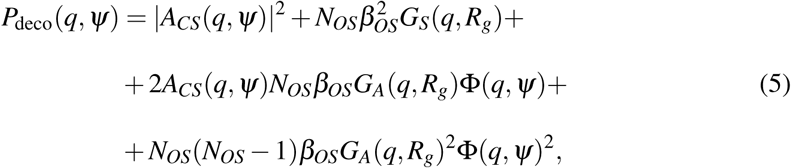

where

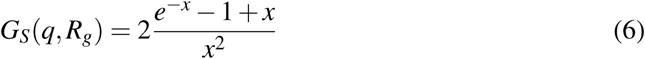

is the structure factor of a Gaussian chain and

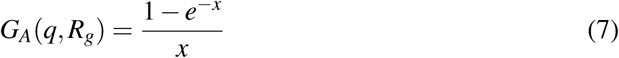

its scattering amplitude, with *x* = (*qR*_*g*_)^2^, where *R*_*g*_ is the radius of gyration of the OS-core. The term Φ(*q, Ψ*) in Eq. 5 is related to the “cross-term” resulting from a uniform distribution of OS-cores all along the ellipsoidal surface of a single bacterium (Pedersen, 2000) and is given by:

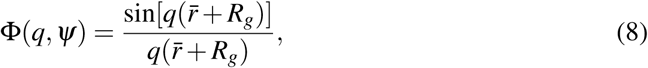

where, according to Eq. 4, 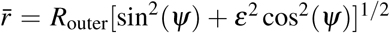. Here, *R*_outer_ is the radius describing the external surface of the cell, which is a function of the *R*_1_ + Δ_*i*_ values in Eq. 2.

The total scattering intensity for a suspension of live *E. coli* cells of number density *n* then reads as:

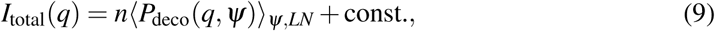

where 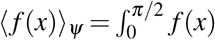 sin(*Ψ*)*dΨ* is the orientational average and 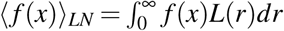 describes the polydispersity of the thickness of the periplasmic space. Specifically, we applied a log-normal distribution function *L*(*r*). The additive constant in Eq. 9 takes into account scattering background at high-*q* originating from unidentified contributions. The log-normal distribution of the periplasmic thickness takes care of the lower cut-off in intermembrane distance fluctuations, given by the finite size of the cell-wall architecture. Note that cell size variations were not considered due to overparameterization, indeed, the polydispersity over the cell radius brought about insignificant changes in the middle to high *q*-range, i.e. *q ≥* 0.05 nm^*−*1^.

## 4. Considering a heterogeneously structured cytosol

It is legitimate to question whether the internal cytosolic structure can be resolved at least in part by SAXS/SANS. In order to address this issue, we derived a more complex analytical scattering function that can be tested against experimental data.

Let us start by considering the general case of a sphere (radius: *R*_0_, SLD: *ρ*_0_), suspended in a medium of *ρ*_*M*_, containing *N* smaller identical, spherical beads (*R*_1_, *ρ*_1_), where the relative distance between two beads *r*_*i*_ *< R*_0_ *−R*_1_. The scattering amplitude of this system is:

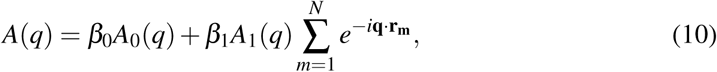

where *β*_0_ = (*ρ*_0_ *− ρ*_*M*_)*V*_0_ and *β*_1_ = (*ρ*_1_ *− ρ*_0_)*V*_1_, and *V*_0_, *V*_1_, *A*_0_ and *A*_1_ are, respectively, the volumes of the sphere and a single bead, and the scattering amplitude of the sphere and a bead. The vector **r**_**m**_ defines all relative distances between the internal beads. The total form factor (*P*(*q*) = |*A*(*q*)|2) then reads as:

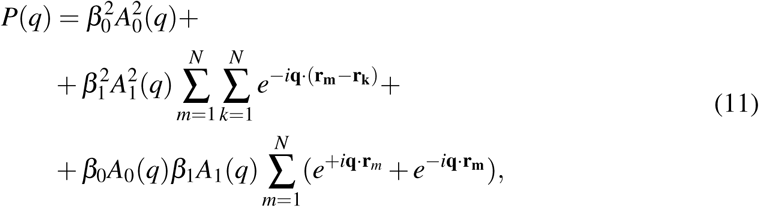

where the first term is the form factor of the sphere; the second term is the total scattering intensity of the *N* internal beads; and the third describes the cross term between the sphere and the beads. Assuming *R*_1_ *≪ R*_0_ and *N →* ∞, one can approximately describe the interior of this sphere as a macroscopic canonical system. This enables the application of the canonical *ensemble* average *(…)*_*N*_ to the summations of Eq. 11 (Klein & D’Aguanno, 1996), leading to:

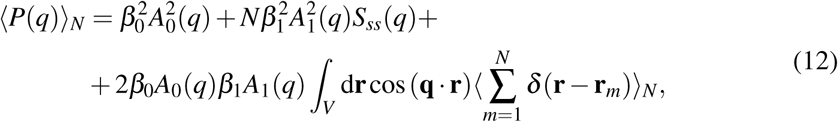

where *S*_*ss*_(*q*) corresponds to the structure factor of interactions among the beads. The summation within the integral defines the microscopic density of *N* beads. Its *ensemble* average corresponds to the single-particle density, which, due to the translational invariance of a homogeneous system, is equal to average bead density *N/V*_0_ (Klein & D’Aguanno, 1996), yielding the final form of sphere-beads system form factor:

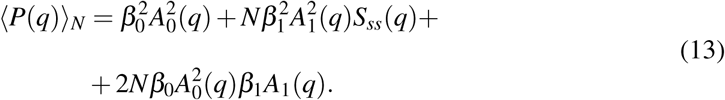

The cross-term is thus modulated by the scattering amplitude of the sphere, *A*_0_, which is equivalent to a convolution of a homogeneous distribution of beads within the volume of a sphere of radius *R*_0_. Importantly, this final form of the cross-term does not depend on the original volume of the sphere hosting the small spheres. That is, even if the beads are constrained within a specific region of the sphere – such as in the case of ribosomes, which mainly partition into the non-nucleoid region of the cytoplasm – the scaling of this cross term does not change.

In the next step, we translate the sphere-bead system to the case of a heterogeneously structured bacterial cytosol. This requires a couple of approximations. First of all, encouraged by the fact that the scattering intensity scales proportionally to the square of the particle volume, we focus on contributions originating from ribosomes. More specifically, ribosomes with a volume *V*_*rb*_ *∼* 2800 nm^3^, a total number *N*_*rb*_ *∼* (1 *−* 6) *×* 10^4^ and *R*_*g*_ = 8.77 nm (Zimmerman & Trach, 1991; Lebedev *et al*., 2015) should be the dominating cytosolic scattering in the case of X-rays or neutrons in the absence of heavy water (Fig. 2 and SI, for cytoplasmic composition and volume fractions). Secondly, we approximate the scattering of ribosomes at very low *q*, i.e. in the Guinier regime up to the first minimum of the scattering form factor, by the scattering of an ‘effective’ sphere of *V*_*rb*_ = 2800 nm^3^ with *R*_*rb*_ = 11 nm. Thirdly, we assume that mixed interactions with macromolecules of different sizes (cytosolic proteins) lead to *S*_*ss*_(*q*) *∼* 1. Further, we simplify the cross-term modulation by a compact ellipsoid, neglecting that ribosomes sequester to the non-nucleotide region, and do not include the grafted OS-cores in the cross-term.

Then the final form of the scattered intensity, considering contributions from ribosomes is given by:

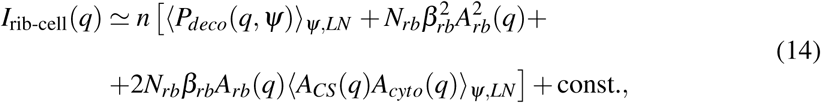

where *β*_*rb*_ = *V*_*rb*_(*ρ*_*rb*_ *−ρ*_*cyto*_), *A*_*rb*_ is the scattering amplitude of ribosomes approximated by an equivalent sphere, and *A*_*cyto*_ is the scattering amplitude of a prolate describing the cytoplasmic space (i.e. only the central part of the core-shell system defining *A*_*CS*_).

## 5. Parametrization and optimization strategy

All parameters needed to describe elastic SAS from *E. coli* are summarized in Tab. 1. In general we differentiate between parameters that either do or do not depend on the individual experiment (e.g. sample concentration, scattering contrast). Experiment-specific parameters were termed “local”, while others were designated as “global” parameters.

**Table 1.** Overview of parameters of the revised multi-scale model for live E. coli scattering (cf. Eqs. 9 and 14).

|  | Parameters | Description |
| --- | --- | --- |
| **Cell body** | <sup>b</sup> $n$ (ml<sup>-1</sup>) | Cell number density |
| | <sup>a</sup> $R$ (nm) | Cell radius, centered at the center of mass of CM |
| | <sup>b</sup> $\rho_{CP}$ (nm<sup>-2</sup>) | Average SLD of the CP core |
| | <sup>a</sup> $\epsilon$ | Ratio between major and minor radii |
| **Ultrastructure profile** | <sup>a</sup> $D_{CM}$ (nm) | Center-to-center distance between the head-group layers in CM |
| | <sup>a</sup> $\Delta_{OM}$ (nm) | Center-to-center distance between CM and OM |
| | <sup>a</sup> $\sigma_{OM}$ (nm) | Standard deviation associated to $\Delta_{OM}$ |
| | <sup>a</sup> $D_{OM}$ (nm) | Center-to-center distance between the head-group layers in OM |
| | <sup>a</sup> $\Delta_{PG}$ (nm) | Center-to-center distance between the PG layer and the OM |
| | <sup>a</sup> $W_{ME}$ (nm) | Width of the head-group layers for both CM and OM |
| | <sup>a</sup> $W_{PG}$ (nm) | Width of the PG layer |
| | <sup>b</sup> $\rho_{TI}$ (nm<sup>-2</sup>) | Average SLD of the tail-group layer in the CM |
| | <sup>b</sup> $\rho_{TO}$ (nm<sup>-2</sup>) | Average SLD of the tail-group layer in the OM |
| | <sup>b</sup> $\rho_{PP}$ (nm<sup>-2</sup>) | Average SLD of the PP layer |
| | <sup>b</sup> $\rho_{ME}$ (nm<sup>-2</sup>) | Average SLD of both CM and OM head-group layers |
| | <sup>b</sup> $\rho_{PG}$ (nm<sup>-2</sup>) | Average SLD of the PG layer |
| | <sup>b</sup> $\rho_{BF}$ (nm<sup>-2</sup>) | SLD of the buffer solution |
| **OS-grafting** | <sup>a</sup> $N_{OS}$ | Number of OS-cores, i.e. LPS molecules |
| | <sup>b</sup> $\beta_{OS}$ (nm) | $\beta$ value for each OS-core |
| | <sup>a</sup> $R_{g,OS}$ (nm) | Effective radius of gyration of each OS-core |
| **Ribosomes** | <sup>c</sup> $N_{rb}$ | Number of ribosomes per cell |
| | <sup>c</sup> $V_{rb}$ (nm<sup>3</sup>) | Volume of a ribosome |
| | <sup>c</sup> $\rho_{rb}$ (nm<sup>-2</sup>) | SLD of a ribosome |
| | <sup>c</sup> $R_{rb}$ (nm) | Radius of the “effective” sphere describing a ribosome |

(a) Global parameter. (b) Local parameter. (c) Mixed global and local, but not tested against SANS data. For details, see main text.

Further, parameters describing structural details of the cytoplasmic and outer membranes were fixed according to values reported from experiments and simulations on membrane mimic systems of *E. coli* cell membranes (De Siervo, 1969; Oursel *et al*., 2007; Lohner *et al*., 2008; Pandit & Klauda, 2012; Kučerka *et al*., 2012; Kučerka *et al*., 2015; Leber *et al*., 2018), as detailed Tab. 2 (see also Fig. 1 and the SI). This decision can be rationalized by the lack of distinct scattering features for *q >* 0.27 nm^*−*1^, corresponding to distances of *∼* 20 nm. Consequently, structural features *≤* 20 nm are, although contributing to overall scattering, difficult to resolve with appropriate accuracy. Note that membrane proteins are treated similarly to proteins in other compartments as bodies adding individually to the scattered intensity and thus do not contribute to the average SLDs of the inner and outer membranes. Compared to the overall scattering arising from the cell body, their overall contribution can be shown to be negligible. Similarly, the width of the peptidoglycan layer *W*_*PG*_, as well as the radius of gyration of the OS core, *R*_*g,OS*_, were fixed to the values reported in Tab. 2 after analysing their contributions. *W*_*PG*_ values of *∼* 6 nm were reported (Matias *et al*., 2003), and simulations in the range 2.5 *−* 7.5 nm (Labischinski *et al*., 1991) led to insignificant variations of *ρ*_*PG*_. Further, our estimates show that *R*_*g,OS*_ < 1 nm (see SI). However, variations of its value within this constraint do not lead to significant changes within our scattering model, because of *R*_outer_ *» R*_*g,OS*_.

**Table 2.** List of fixed parameter values.

| Parameters | Values |
| --- | --- |
| $D_{CM}$ (nm) | 3.73 |
| $D_{OM}$ (nm) | 3.33 |
| $W_{ME}$ (nm) | 0.75 |
| $\rho_{TI} \times 10^{-4}$ (nm$^{-2}$) | 8.31<sup>a</sup> / 0.022<sup>b</sup> |
| $\rho_{TO} \times 10^{-4}$ (nm$^{-2}$) | 8.86<sup>a</sup> / 0.012<sup>b</sup> |
| $\rho_{ME} \times 10^{-4}$ (nm$^{-2}$) | 12.9<sup>a</sup> / 1.24–4.11<sup>b,c</sup> |
| $\varepsilon$ | 2.0/1.75<sup>d</sup> |
| $W_{PG}$ (nm) | 6.0 |
| $R_{g,OS}$ (nm) | 0.45 |

(a) X-ray SLDs. (b) Neutron SLDs. (c) These two values respectively refer to hydrated head-groups SLDs with 0 and 100 wt% D<sub>2</sub>O buffer compositions, accounting for exchangeable hydrogens. (d) $\varepsilon = 2$ for ATCC 25922 and 1.75 for K12 5K strains.

Finally, the ratio between major and minor ellipsoidal radii, *ε*, was fixed for ATCC and K12 strains. This choice was driven by the lowest available *q*-range, which does not reach the Guinier plateau of the bacterial scattering and thus does not allow for an accurate determination of the cell length. We therefore used dynamic light scattering to first estimate the average length, *L*_*c*_, from 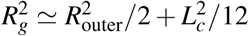, using *R*_*g*_ *≈ R*_*H*_ and typical values reported for *R*_outer_ in literature. This led to 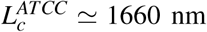 and 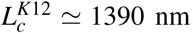. The resulting *ε* = *L*_*c*_*/*(2*R*_outer_) values were subsequently refined in some test runs of the optimization procedure for USAXS/SAXS data with *R*_outer_ as adjustable parameter and then fixed for the detailed USAXS/SAXS and VSANS/SANS analysis, yielding the values reported in Tab. 2.

Due to the complexity of the system and the high number of parameters, optimization of the adjustable parameters was performed with a Monte Carlo genetic selection algorithm (Banzhaf *et al*., 1998). In brief, the algorithm is schematized in a series of steps that are repeated at every cycle (*generation*). As step zero, nine low-discrepancy sequences (quasirandom numbers) were created for each parameter (*gene*) within specific boundaries, based on compositional estimates (see the SI). Nine sets of parameters (*individuals*) were then used as input for testing an equal number of possible scattering intensity curves (step 1, *evaluation*). This involved the calculation of nine standard weighted chi-squared values as:

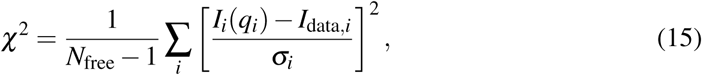

where *N*_free_ is the number of free parameters, and *σ*_*i*_ is the error associated to the measured *I*_data,*i*_ at a given *q*_*i*_. Only the four *individuals* with the lowest *χ*2-values were then selected (step 2, *selection*) to generate the first *offspring*, consisting of eight new *individuals* that were created by randomly shuffling the *genes* of the selected four parents (step 3 *recombination*). The new 9th *individual* instead was a copy of the parent with the lowest *χ*2-value (rule 1: *inheritance of the best*). After the *recombination*, each *gene* had a finite probability to be altered (step 4, *mutation*). In addition, there was also a finite probability of replacing an entire *individual* with a brand new set of randomly created *genes* (rule 2: *the stranger*). This construction of the new *offspring* ended the first cycle, and each *individual* was again evaluated on the base of the *χ*2 values (jumping back to step 1). The process was repeated until the changes of the lowest 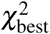 were less than 0.5% for 25 consecutive *generations*.

The *mutation* step and rule 2 allow the algorithm to skip possible local minima in the *χ*2 landscape, whereas the rule 1 enables a fast convergence of the fitting. Note that, except for the creation of the initial set of *individuals*, pseudorandom numbers were used in the whole algorithm for the decision-making processes of *recombination* and *mutation* steps, and for rule 2. Note also that a bigger initial population (*>* 9 *individuals*) did not result in a gain of computational time. Each new pseudorandom *gene* was always constrained within the initial boundaries, in order to assure the preservation of a physical meaning of the results. In total, each scattering curve was fitted 500 times, and only converging fittings (convergence criterion 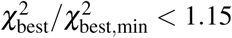 were used to retrieve average values and standard deviations for each parameter.

In the case of SANS, instrumental smearing was additionally accounted for by convoluting the calculated scattered intensities with a Gaussian function of width Δ*q*(*q*), using values as provided by the D11 primary data treatment.

## 6. Results and discussion

### 6.1. SAXS/SANS global analysis

The revised multi-scale model was tested against USAXS/SAXS and VSANS/SANS data on *E. coli* strains ATCC 25922. Ten differently contrasted SANS data-sets were collected, varying D_2_O from 0 wt% to 90 wt% (increments of 10 wt%). Results of the combined SAXS/SANS data analysis are reported in Fig. 3 and Tabs. 3 and S1. Fig. 3A specifically highlights the different contributions from the multi-core-shell model, the OS-cores and the sum of the two cross-terms (see Eq. 5) in the USAXS and SAXS regimes. The scattering contribution from the core-shell function, i.e. cell-body plus cell-wall, dominates over the scattering intensity originating from the OS cores, due to the huge difference in mass. However, the cross term, being a function of the whole cell surface is mainly responsible for modulating the scattered intensities between *q ∼* 0.1 and 0.3 nm^*−*1^. This leads to an average slope between *q*^*−*1.5^ and *q*^*−*2^ in this regime, which is a typical signature of grafted systems, also called “blob scattering” (Pedersen, 2000). Previously, this regime was taken to be dominated by flagella, described as a self-avoiding-walk polymer term (Semeraro *et al*., 2017). This however does not describe the apparent change of slope at *q ∼* 0.04 nm^*−*1^ and the scattering feature at *q ∼* 0.1 nm^*−*1^, which appears to be specific to the ATCC strain (see Fig. S3), but not K12 (see below). Hence, the OS-core cross-term enables a full description of the *q*-range between 0.03 and 0.2 nm^*−*1^.

**Table 3.** Fit results for the global parameters describing USAXS/SAXS and VSANS/VANS of E. coli ATCC 25922 strain. Errors were calculated from standard deviations of the ensemble of converged fittings. See Tab. S1 for results on local parameters.

|  | USAXS/SAXS | VSANS/SANS |
| --- | --- | --- |
| $R$ | $371 \pm 3$ nm | $369 \pm 3$ nm |
| $\epsilon$ | $2.0^a$ | |
| $\Delta_{OM}$ (nm) | $34.3 \pm 1.0$ | $32.0 \pm 1.0$ |
| $\sigma_{OM}$ (nm) | $7.4 \pm 0.7$ | $7.8 \pm 0.3$ |
| $\Delta_{PG}$ (nm) | $17.8 \pm 0.2$ | $16.7 \pm 1.7$ |
| $N_{OS}$ | $(4.7 \pm 0.3) \times 10^6$ | $(6.2 \pm 0.6) \times 10^6$ |

(a) Fixed parameter.

**Fig. 3.**
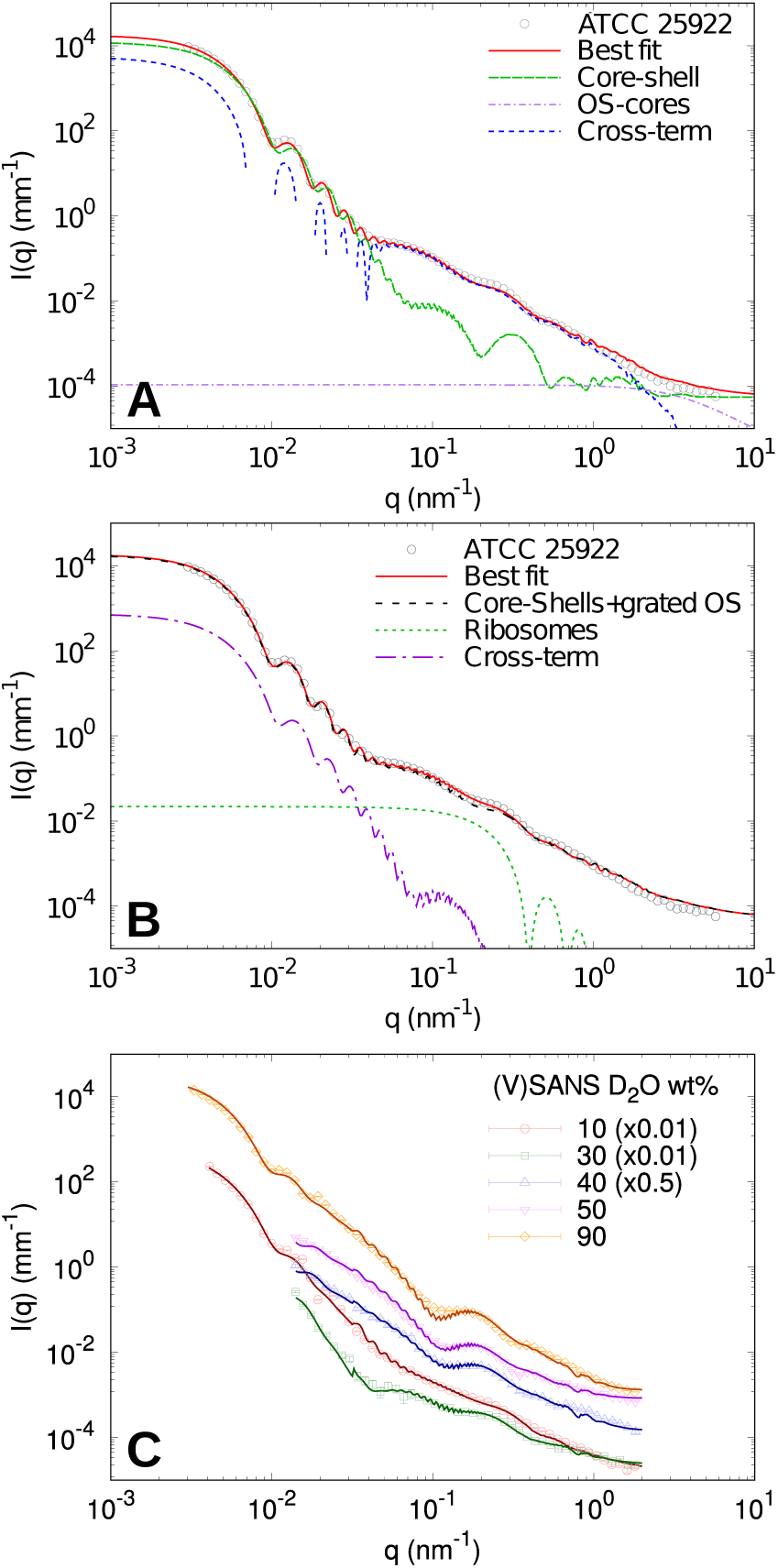
A) USAXS/SAXS data analysis of *E. coli* ATCC 25922 using Eq. (9) highlighting contributions from different terms (negative values of the cross term are not shown). B) Alternative analysis of the same data using Eq. (14), showing contributions from ribosomes. Comparison to a fit using Eq. 9 (black dashed line) shows negligible differences. C) VSANS/SANS data of the same strain at selected D_2_O contrasts (see also Fig. S4 for additional neutron data). Scattering curves were scaled for a better visibility.

Attempts to fit the same data with the more complex model accounting for ribosomes (Eq. 14) demonstrated insignificant scattering contributions from the macromolecules (Fig. 3B). In particular, the goodness-of-fit, as well as results for the common adjustable were identical within the error of the analysis. Thus, the ribosome term, along with its related cross-term, is negligible compared to the cell-wall contribution. Note that only SAXS or SANS data at the lowest wt% of D_2_O are sensitive to test for this contribution. SANS data is dominated by the acyl-chain contribution at higher heavy water content (Fig. 2). Further, ribosomes are made up of amino acids and RNA, which will be differently matched, thus challenging the analysis. Note that *N*_*rb*_, was the only adjustable parameter for this analysis. *V*_*rb*_ and *R*_*rb*_ values were fixed as detailed in section 4. Interestingly, the outcome of this test resulted in a much smaller number of ribosomes (*N*_*rb*_ *∼* 500) than our estimate of *N*_*rb*_ *∼* 10^4^ based on (Zimmerman & Trach, 1991), see the SI. This is possibly related to the low contrast of these molecules in the local cytoplasmic environment. Indeed, as the scaling is proportional to *N*_*rb*_(*ρ*_*rb*_ *− ρ*_*CP*_)^2^, small differences in the effective contrast easily skew the determination of the number of macromolecules.

Importantly, this analysis not only demonstrates that the effective scattering signal from ribosomes is negligible, but similar considerations can also be applied to the other cytoplamic components. However, because of their either smaller size (proteins) or volume fraction (DNA and RNA), they will contribute even less to the overall scattered intensity. Hence, elastic scattering techniques are not suitable for discriminating differently structured compartments within the cytosol in live bacterial cells. The same applies also to membrane proteins (see above) or proteins present in the periplasmic space and peptidoglycan layer.

Analysis of selected VSANS/SANS data at selected contrasts is presented in Fig. 3C (the entire set of neutron scattering data and fits is presented in Fig. S4). Clearly, fits using Eq. (9) neatly capture all changes of scattered intensities upon varying D_2_O concentrations, lending strong support to our modelling approach. The resulting parameters forming the X-ray and neutron SLD profiles of the bacterial envelope are summarized in Fig. 4, again at selected neutron contrasts (see, Fig. S5, for all neutron SLD profiles). Small differences between distances in X-ray and neutron profiles, as well as global parameters (Tab. 3), are due to biological variability of the samples, but are, with the exception of *N*_*OS*_, still within the confidence range of the results. At 10 wt% D_2_O, the contrast differences between different slabs are comparable with those obtained from SAXS data. Indeed, the scattering intensities in these cases were also comparable in terms of scattering features at *q* = 0.1 and 0.3 nm^*−*1^ (Fig. 3). In turn, at 40 D_2_O wt% (and similarly up to 90 wt% D_2_O), the contrast of highly hydrated bacterial subcompartments (PG layer, etc.) is much lower than the major contrast of the hydrophobic regions of the two membranes (Fig. 4B). This characteristic leads to the shift in the scattering feature from *q ∼* 0.27 to *q ∼* 0.2 nm^*−*1^ (Fig. 3C), which is primarily related to the intermembrane distance. This is in good qualitative agreement with the invariant estimation (Fig. 2), which suggests that the scattering intensity is dominated by the contribution from the acyl chain region for D_2_O *≥* 40 wt%.

**Fig. 4.**
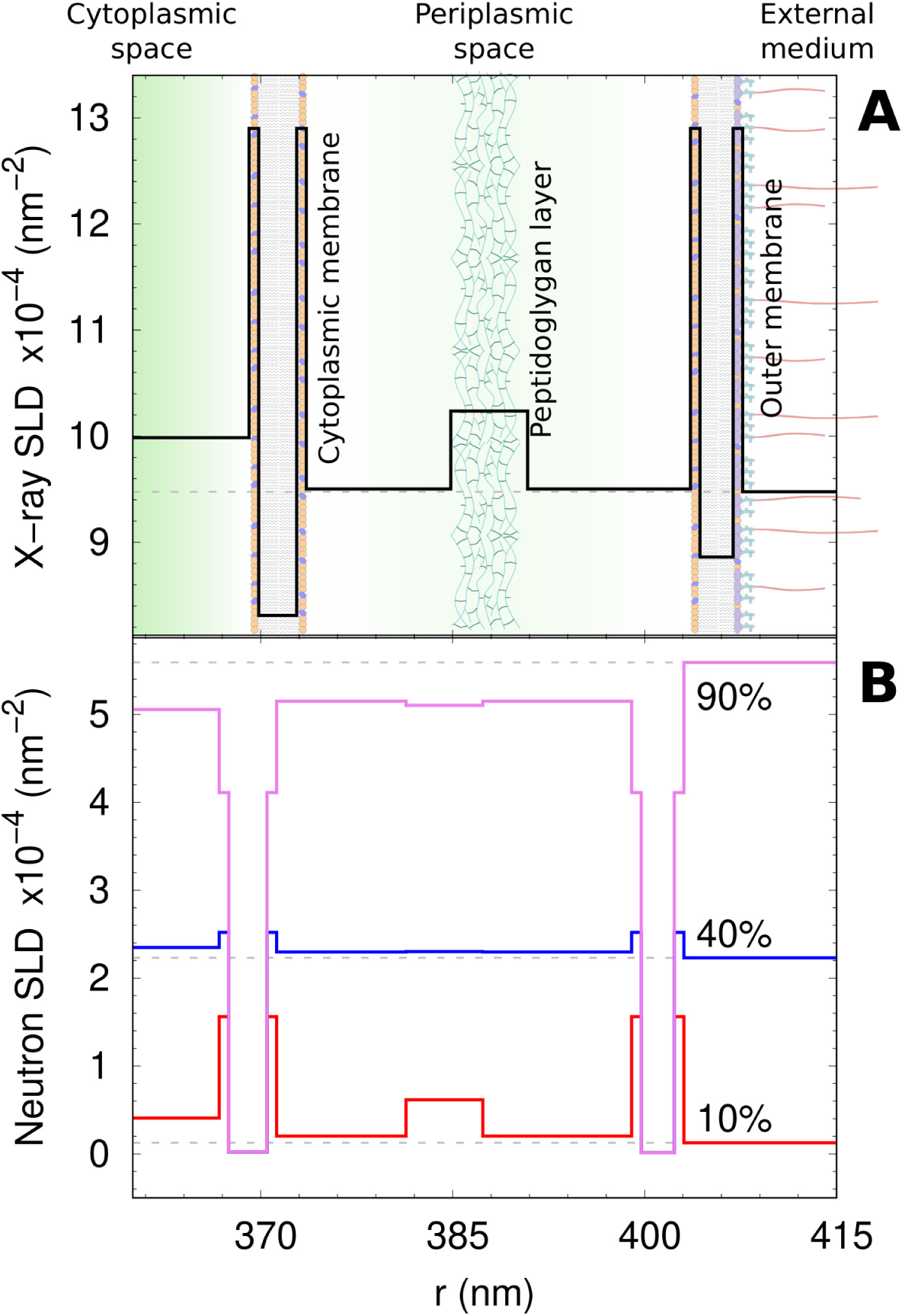
A) X-ray SLD profile of the bacterial ultrastructure of ATCC 25922 strain, corresponding to the fit shown in Fig. 3A. The panel highlights the average positions of both cytoplasmic and outer membrane, and the peptidoglycan layer. The abscissa describes the distance from the cell center along the minor radius *R*. B) Selected neutron SLD profiles of the same strain (cf. Fig. 3C). See also Tab. S1

As a final sanity check for our modelling, we report the variation of the various SLDs with D_2_O content. Since the solvent freely accesses the cytoplasmic and periplasmic spaces, such plots should display a linear dependency. Indeed, the trends followed the expected behavior, which also enabled us to calculate the match points for the individual compartments (Fig. 5 A-C). Note that the SLD values of the hydrated phospholipid headgroups were fixed (Tab. 2). In the case of *β*_*OS*_, scattering contributions are superseded by the signal originating from the intermembrane distance for D_2_O *<* 60 wt%. The linear trend for *β*_*OS*_ was therefore determined in the range 0 *−* 50 wt%, and then extrapolated to higher D_2_O concentrations by using a confidence boundary of *±*20%. Results from this analysis were used to derive the measured effective invariant as a function of D_2_O wt% (Fig. 5D). The comparison with the estimated *Q* shows a shift in the minimum from the estimated 40 wt% to the measured 50 D_2_O wt%, possibly due to a bigger contribution from the components that dominate at D_2_O *≤* 30 wt% (Fig. 2). On the other hand, these components are the very macromolecules that were proven to have a negligible scattering contribution.

**Fig. 5.**
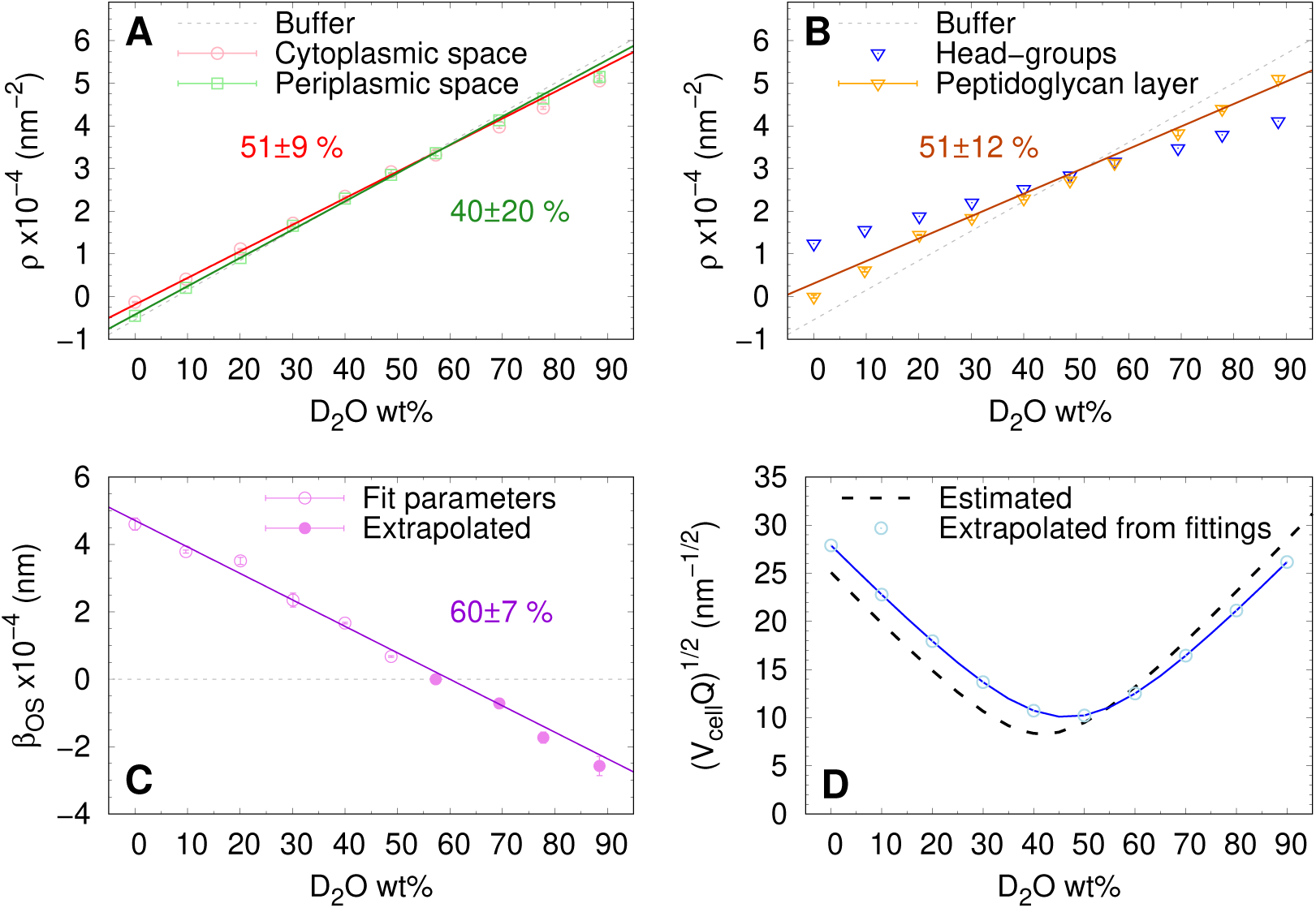
A-B) Plot of the cytoplasm (red circles), periplasm (green squares) and peptidoglycan (orange triangles) SLDs, along with linear fittings and matching points. The SLDs of the phospholipid head-group layers were fixed parameters (blue triangles). C) Plot of the *β*_*OS*_ (purple diamonds) values, along with linear fits and matching points. D) Comparison between estimated scattering invariant and extrapolated forward scattering.

The answer to this paradox can be found in the absence of a net size distinction between macromolecules that scatter individually or are included in SLD averages. Translating the molecular mass distribution from gel filtration of cytosolic proteins by (Zimmerman & Trach, 1991) into a size probability-distribution-function, yields a maximum value at the smallest protein size, detected in our experiments (see SI). Hence, it is likely that a fraction of the smallest bacterial proteins need to be included in the parameter *ρ*_*CP*_. Our fitted values for *ρ*_*CP*_ are indeed larger than the estimated SLDs of the metabolites in the case of both SAXS and SANS analysis (compare Fig. 1 and Tab. S1). In any case, the estimated invarant was used as a valuable guide for our modelling, only. Obviously, Eq. 1 loses its validity in the case of dense and crowded suspensions, and a new formula accounting for the each volume fraction should be used for improved estimates (Porod, 1982).

### 6.2. Comparison between ATCC and K12-related strains

After successful verification of our revised multi-scale model for live *E. coli* ATCC, we tested whether the new model is also applicable to other *E. coli* strains. Fig. 6 shows the USAXS/SAXS data of the K12 5K, JW4283 and Nissle 1917 strains in comparison to ATCC. Strikingly, the scattering patterns of K12 5K, JW4283 (which is fimbriae-free) and Nissle 1917 are superimposable for *q >* 0.06 nm^*−*1^, suggesting that the main ultrastructural features are conserved in these strains and confirming that the presence of fimbriae does not contribute to SAXS. Note also that our previous model would perfectly fit all K12 strains. Different minima positions of the scattered intensities at lower *q*-values are due to the different sizes of the different strains, instead.

**Fig. 6.**
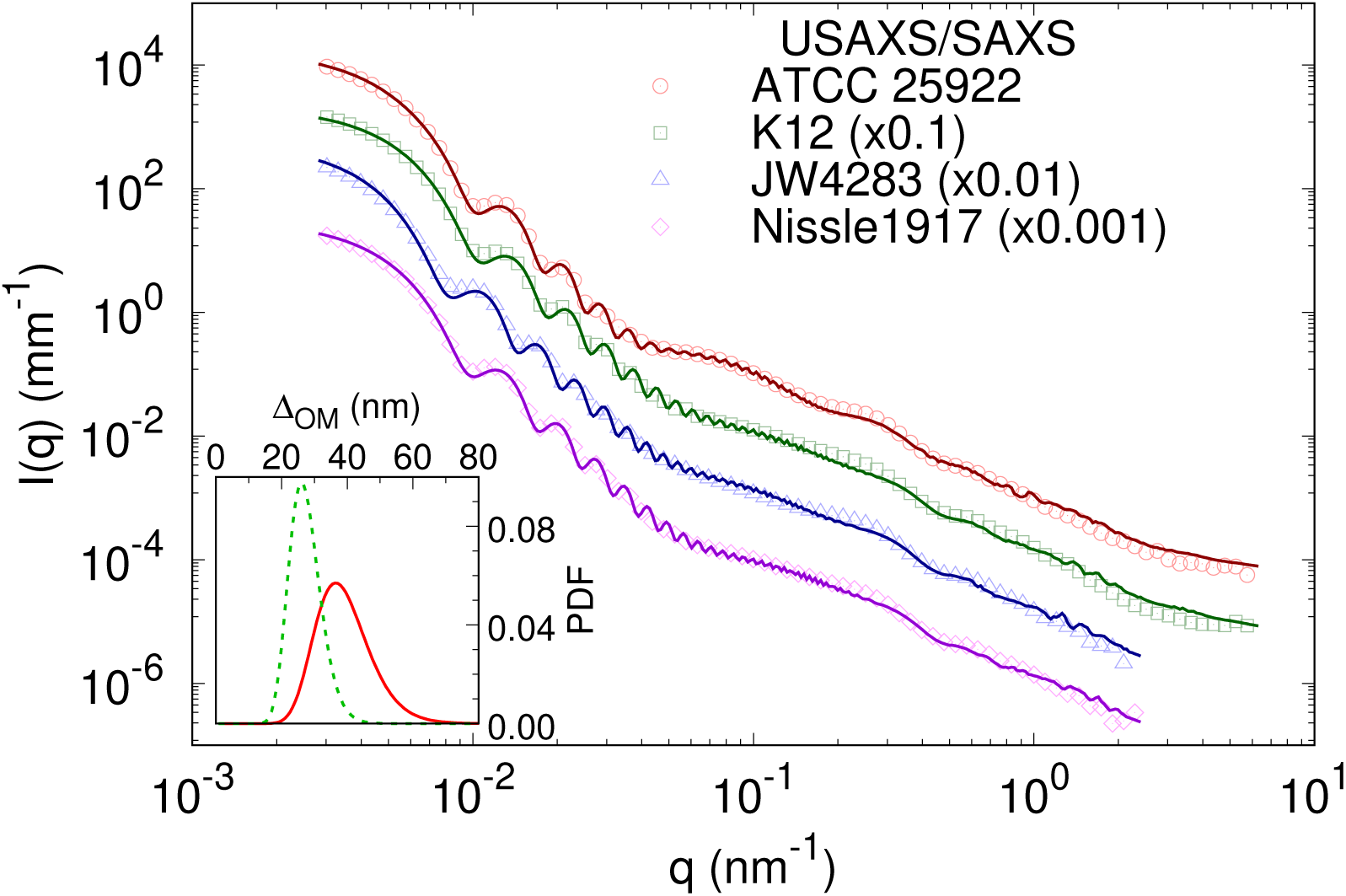
Multi-scale analysis of USAXS/SAXS data of the ATCC, K12, fimbria-free K12 JW4283 and Nissle 1917 strains. The inset shows the plots of the log-normal probability distribution function (PDF) of Δ_*OM*_ values for ATCC (solid red) and K12 (dashed green) strains. The PDFs of JW4283 and Nissle 1917 are comparable to K12 and are thus not shown.

Importantly, our new model is capable of fitting all strains, as demonstrated by the overall excellent agreement with experimental data (Fig. 6). Structural parameters resulting from this analysis are reported in Tab. 4 and Tab. S2. Most of these parameters are of comparable magnitude. Significant differences concern the cell size (*R, ε*) – as observed in the different positions of the scattering minima (Fig. 6), the number of OS-cores (*N*_*OS*_) and the intermembrane distace (Δ_*OM*_, *σ*_*OM*_). The latter is related to the actual periplasmic space thickness via Δ_*OM*_ *−* (2*W*_*ME*_ + *D*_*CM*_ + *D*_*OM*_)*/*2 (see, Fig. S6, for the X-ray SLD profiles). Both, periplasmic thickness and its fluctuation are smaller for K12-related strains than for ATCC. Note that Δ_*OM*_ for K12 5K is consistent with our previously reported value for a similar strain (Semeraro *et al*., 2017). Despite the different Δ_*OM*_ and *σ*_*OM*_ for either ATCC or K12 strains, the magnitude of the relative fluctuations *σ*_*OM*_*/*Δ_*OM*_ *∼* (0.16 *−* 0.22) is roughly conserved.

**Table 4.** Fitting results for the set of local free parameters for USAXS/SAXS analysis of ATCC 25922, K12 5K, JW4283 and Nissle 1917 strains.

| **Free parameters** |  |  |  |  |
| --- | --- | --- | --- | --- |
|  | ATCC 25922 | K12 5K | JW4283 | Nissle 1917 |
| $R$ (nm) | $371 \pm 3$ | $363 \pm 3$ | $471 \pm 4$ | $397 \pm 3$ |
| $\epsilon$ | $2.0^a$ | $1.75^a$ | $1.71 \pm 0.03$ | $1.75^a$ |
| $\Delta_{OM}$ (nm) | $34.3 \pm 1.0$ | $23.8 \pm 0.6$ | $26.5 \pm 0.8$ | $23.4 \pm 0.6$ |
| $\sigma_{OM}$ (nm) | $7.4 \pm 0.3$ | $4.2 \pm 0.2$ | $5.4 \pm 0.3$ | $3.7 \pm 0.2$ |
| $\Delta_{PG}$ (nm) | $17.8 \pm 0.2$ | $16.8 \pm 0.2$ | $17.8 \pm 0.2$ | $17.3 \pm 0.2$ |
| $N_{OS} (\times 10^6)$ | $4.7 \pm 0.3$ | $4.10 \pm 0.12$ | $6.0 \pm 0.6$ | $4.09 \pm 0.17$ |

(a) Fixed value.

Considering cell size differences we find, based on the cell surface, an order that follows K12 5K (2.8 *×* 10^6^ nm^2^) *<* Nissle 1917 (3.3 *×* 10^6^ nm^2^) *≈* ATCC 25922 (3.4 *×* 10^6^ nm^2^) *<* JW4283 (4.5 *×* 10^6^ nm^2^). Differences in cell size are expected to be coupled to the number of LPS molecules, dominating the outer leaflet of the cellular envelope. Indeed, *N*_*OS*_, follows roughly the order observed for the bacterial outer surface (Tab. 4). Normalizing *N*_*OS*_-values by the bacterial outer surface leads to a LPS surface density of 1.3 *−* 1.5 nm^*−*2^. However, as the cross-sectional area per LPS is *∼* 1.6 nm^2^ (Clifton *et al*., 2013; Micciulla *et al*., 2019; Kim *et al*., 2016), the expected surface density is *∼* 0.6 nm^*−*2^. The discrepancy between the two estimates is most likely due to an underestimation of the bacterial surface by considering the prolate approximation, or uncertainties introduced by *β*_*OS*_, which, as *N*_*OS*_, scales scattering contributions from oligosaccharides and their cross-terms (Eq. 5). An additional factor could be related to the roughness of the bacterial surface (Alves *et al*., 2010), which results in a larger effective surface than considered here in our simple estimate.

Finally, the center-to-center distance between the PG layer and the OM, Δ_*PG*_ *∼* 17 nm, with a X-ray SLD *ρ*_*PG*_ *∼* 10.2 *×* 10^*−*4^ nm^*−*1^ for all presently studied *E. coli* strains. Previously, we reported Δ_*PG*_ *∼* 11 nm (Semeraro *et al*., 2017), which appears to be more consistent with the length of the lipoproteins cross-linking the peptidoglycan strands to the outer membrane. This deviation from the expected value might be due to the fluctuation modes of Δ_*PG*_ that are not fully correlated to those of Δ_*OM*_, which here is modelled by a log-normal distribution function. Devising a separate/partially-coupled distribution function for variations of Δ_*PG*_, is beyond the present experimental resolution, however. In contrast, our new value for 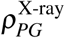 (and also 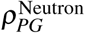 for ATCC) is now consistent with reported hydration values of the peptidoglycan layer, i.e. 80-90 vol% (Labischinski *et al*., 1991; Pink *et al*., 2000). The previously reported value *∼* 11.6 *×* 10^*−*4^ nm^*−*1^ included the presence of macromolecular species in the SLD average.

## Conclusion

The similarity of SAXS data of native and flagella/fimbriae-free *E. coli* strains led us to revise our previously reported scattering form factor model (Semeraro *et al*., 2017) of the Gram-negative bacterium *E. coli*. The flagellar contribution was substituted by considering the scattering from the oligosaccharide inner and outer cores of the lipopolysaccharides, in terms of a grafted-polymer model. The model presented here is based on detailed compositional and structural estimates of characteristic lengths, volumes and scattering length densities for each cellular component and thus unifies decades of research on *E. coli* ultrastructure and molecular composition into a single comprehensive scattering function. The applicability of the derived model to X-ray and neutron scattering experiments enables the use of the powerful technique of contrast variation in order to highlight or nullify contributions from specific bacterial compartments.

Interestingly, we found that combined (U)SAXS/(V)SANS experiments are not sensitive to the structural heterogeneity of the cytoplasm as the scattering signal of its constituent macro-molecules is being overwhelmed by the contribution from the cell envelope. Likewise, the combined analysis is not able to report differences in the sub-nanometer range, in particular for cytoplasmic or outer membranes, such as thickness or compositional asymmetry to name but a few. The underlying SLD variations for CM and OM, were therefore fixed at values detailed in Tab. 2, along with the width of the peptidoglycan layer and the effective *R*_*g*_ of each oligosaccharide core. In turn, our technique is highly sensitive to the overall cellular size, the average contrast of the cytoplasmic and periplasmic space, as well as the structure of the cellular envelope. The latter includes the distance between cytoplasmic and outer membranes, as well as its average fluctuations, the distance between the peptidoglycan layer and the outer membrane, and the overall number of LPS molecules (oligosaccaride cores) in the outer membrane.

Complementary to transmission electron microscopy or optical microscopy, elastic scattering experiments on live *E. coli* therefore provide ensemble averaged values of specific ultrastructural bacterial features without the need of invasive labelling. Here we report differences between five different *E. coli* strains, which were mainly due to overall size and intermembrane distances (Tab. 6). Future research may exploit this platform to detect effects of different sample growth conditions or the effects of bactericidal compounds such as antibiotics. Especially the combination of our analysis with millisecond time-resolved (U)SAXS enables kinematographic detection of their activity. Our laboratory is currently exploring such an approach for antimicrobial peptides.

This work was conducted in the framework of the Austrian Science Fund (FWF) project No. P30921 (awarded to KL). ESRF – The European Synchrotron and the Institut Laue-Langevin (ILL) are acknowledged for provision of, respectively, SAXS (proposal LS-2869) and SANS (exp. 8-03-910) beamtimes. The authors would also like to thank the Visitors lab support at EMBL Grenoble for providing access to the lab equipment for bacterial sample preparation during SAXS and SANS experiments. The authors are grateful for the support of T. Narayanan for the USAXS/SAXS measurements, as well as for the fruitful discussion and advices. Finally, the authors would like to thank the staff of the Institute of Molecular Biosciences, ID02-beamline and D11-instrument for the support and availability.

## Synopsis

Structural and compositional information about *E. coli* cells are summarized and translated into an analytical multi-length-scale scattering form factor model of live bacterial suspensions.

