## Supplementary material for "Evolution of the Analytical Scattering Model of Live *Escherichia Coli*"

### 1. Bacterial structure and composition

Overall, *E. coli* are rod-shaped bacteria (length: $1 - 3\ \mu m$, diameter: $0.7 - 1\ \mu m$), with a density of about $(1.08 - 1.16)\ g^{-1}$ (Schwarz-Linek *et al.*, 2015). The following estimates primarily refer to K12 strain in the mid-exponential growth phase. Differences in composition specific to other *E. coli* strains or other phases of the growth cycle are not considered here, but they can be estimated in a similar fashion and in principle also be accounted for by adjusting the parameters of the scattering model.

#### 1.1. Cytoplasmic space

The cytoplasm is the hydrogel-like core of cell ($\sim 75$ vol% water), composed by freely diffusing macromolecules (DNA, RNA, ribosomes, proteins), low-molecular-weight molecules (e.g. ATP, aminoacids) and ions. The cytoplasmic volume is 'virtually' divided into two regions: the nucleoid area, which is defined by the volume spanned by the main DNA ring; and the non-nucleoid volume, where ribosomes – the biggest cytoplasmic macromolecules after the main DNA ring – are free to diffuse.

**Nucleoid Region** The principal component of this area is the main DNA string, which is a folded ring that can even reach an end-to-end length, $L$, of up to 1 mm in the case of long cells at the late stage of their division cycle. The volume spanned by this entangled 'wire' defines the compact nucleoid region, which is dynamic and does not display any specific shape. The DNA double-helix has a base-pair distance of $0.33-0.34$ nm and an average molecular weight of 309.1 Da per base. By dividing $L$ by the base-pair distance, the number of pairs of such a long string amounts to about $3\times10^6$, with a total weight around $1.9\times10^9$ Da. As its density is about 1.7 g ml$^{-1}$, the actual volume, considering $L = 1$ mm, is only $\sim 1.8\times10^6$ nm$^3$ (i.e. $< 0.1$ vol% of the cytoplasmic volume). This small value should not be confused with the apparent total volume of the nucleoid region, which, in typical TEM images, is visible as a bright area occupying about half of the cytoplasmic volume (Hobot *et al.*, 1984; Beveridge, 1999) and hosts also proteins, mRNA and other low-molecular-weight molecules.

**Non-Nucleoid Region** The non-nucleoid region is crowded with ribosomes, tRNA, mRNA, proteins and plasmids. Estimates for cytoplasmic concentrations of ribosomes, proteins and RNA within the cytoplasm can be obtained by comparing the measured dry masses of each bacterial component (Neidhardt *et al.*, 1990) with total protein and RNA mass and their size distribution in *E. coli* (Zimmerman & Trach, 1991). Only about 5 wt% of the mass of these cytoplasmic high-molecular-weight macromolecules consists of tRNA and mRNA, which is equivalent to a cytoplasmic volume fraction of $\sim 1$ vol% with molecular sizes of $R_g \sim 2-3$ nm (tRNA) and $R_g \sim 6-9$ nm (mRNA). In contrast, the mass fraction of ribosomes (34 wt%) equals 6-8 vol% and a size of $R_g = 8.77$ nm has been reported for *E. coli* ribosomes from

SAXS experiments (Lebedev *et al.*, 2015).

Finally, the estimated protein content (60 wt%), amounts up to 13-16 vol%, with an average $R_g \sim 2$ nm (Zimmerman & Trach, 1991). A more detailed look reveals that the protein size distribution decays exponentially from $R_g \sim 1.5$ nm for the largest protein fraction to few macromolecules with $R_g \sim 4$ nm (Zimmerman & Trach, 1991). All together, proteins, ribosomes and RNA, occupy 20-25% of the cytoplasmic volume.

**Metabolite Pool and Ions** The *E. coli* cytoplasm contains $\sim$ 1200 low-weight molecules, including nucleotides, amino-acids, alcohols, etc. (Tweeddale *et al.*, 1998; Maharjan & Ferenci, 2003; Bennett *et al.*, 2009), all listed in the *E. coli* Metabolome Data Base (ECMDB, http://ecmdb.ca/) (Guo *et al.*, 2012). The metabolite pool of K12 in the exponential growth phase, for example, is dominated by glutamic acid ($\leq$ 150 mM), whereas other major components such as e.g. spermidine ($\leq$ 20 mM) or ATP ($\leq$ 10 mM) are much less abundant. The cytosolic content of simple electrolytes can be surmised as $\sim$ 224 mM $K^+$, $\sim$ 70 mM $Na^+$, and $4-18$ mM $Mg^{+2}$, $Ca^{+2}$, $Fe^{+3}$, and a few more heavy ions. The 35 most abundant metabolites, and the 10 most abundant ionic species were selected from the ECMDB in order to estimate SLDs of the cytoplasmic medium, yielding $9.7 \times 10^{-4}$ nm$^{-2}$, in the case of X-ray SLD, and neutron SLD values of $-0.44 \times 10^{-4}$ nm$^{-2}$ and $6.25 \times 10^{-4}$ nm$^{-2}$, where the latter includes 100 wt% $D_2O$ PBS buffer and full interfacial H/D exchange.

#### *1.2. Cell envelope*

**Cyoplasmic Membrane Lipids** The inner, or cytoplasmic, membrane of *E. coli* is composed by a variety of phospholipids (De Siervo, 1969; Oursel *et al.*, 2007) with the most abundant species being phosphatidylethanolamine (PE), phosphatidylglygerol (PG) and cardiolipin (85:5:10 mol/mol/mol) (Lohner *et al.*, 2008). The fatty acid composition dominated by C16:0 and C18:1 hydrocarbons (De Siervo, 1969). For estimating its SLD it is legitimate to approximate the lipid composition by palmitoyl-oleoyl-PE (POPE) and palmitoyl-oleoyl-

PG (POPG) (3:1 mol/mol) (Leber *et al.*, 2018). This approximation allows us to use previously reported high-resolution structural data of pure POPG and POPE bilayers (Kučerka *et al.*, 2012; Kučerka *et al.*, 2015), yielding $\rho_{TI} = 8.31 \times 10^{-4}$ nm$^{-2}$ (X-rays) and $\rho_{TI} = 0.022 \times 10^{-4}$ nm$^{-2}$ (neutrons) for the acyl chain region. In order to estimate the average SLDs for the lipid head-groups, we assumed a hydration $\sim 40$ vol%, leading to $\rho_{ME}^{\text{X-ray}} \sim 12.9 \times 10^{-4}$ nm$^{-2}$, and $\rho_{ME}^{\text{neutron}} \sim 1.24 \times 10^{-4}$ nm$^{-2}$ and $4.43 \times 10^{-4}$ nm$^{-2}$, respectively, where the latter value includes 100 wt% $D_2O$ PBS buffer and full interfacial H/D exchange of the PE and PG groups. A molecular dynamics simulation of a realistic *E. coli* K12 lipid mixture (Pandit & Klauda, 2012) was taken as reference to estimate the membrane thickness. This accounts for the variety of acyl chain lengths and gave 2.98 nm for the hydrophobic length, and 3.73 nm for the average bilayer thickness.

**Outer Membrane Lipids** While the composition of the inner leaflet of the outer membranes closely resembles that of the inner membrane, its outer leaflet is almost exclusively composed of lipopolysaccharides (LPS) (Seltmann & Holst, 2002). A single LPS molecule can be decomposed in the lipid A – usually a phosphorylated glucosamine disaccharide decorated with 4 to 7 C14:0 chains (Kim *et al.*, 2016) – the inner and outer oligosaccharide cores, and the O-antigen. The inner and outer cores, include keto-deoxyoctulosonate (kdo), heptose, phosphorylated heptose, and hexose such as glucose or galactose. Especially in the case of the inner core, its composition is well-conserved among strains (Heinrichs *et al.*, 1998; Müller-Loennies *et al.*, 2003). Finally, the O-antigen is a long polysaccharide chain of different sugars (Seltmann & Holst, 2002; Rodriguez-Loureiro *et al.*, 2018). Composition and length of O-antigen chains, as well as their distribution along the surface are very diverse (Kučerka *et al.*, 2008; Micciulla *et al.*, 2019). LPS molecules possessing the O-antigen are called “smooth” LPSs, as in the case of the *E. coli* strain ATCC 25922. Instead, other strains, as the most studied the *E. coli* K12, have “rough” LPSs, i.e. lacking the polysaccharide chains but conserving the OS-cores (Heinrichs *et al.*, 1998).

Estimations of SLDs and lengths characterizing LPS molecules can be also separated into these three building blocks. The SLD from the head-group of lipids A can be approximated as similar to phospholipids, whereas its short acyl chains will have values close to that of myristic acid, $9.4 \times 10^{-4}$ nm$^{-2}$ (X-ray SLD) and $0.0011 \times 10^{-4}$ nm$^{-2}$ (neutron SLD). Similarly, the hydrophobic length and the total membrane thickness (cores excluded) were set to 2.58 nm and 3.33 nm, respectively. The OS core, a branched chain of 13 saccharides (7 in the inner and 6 in the outer core) in the case of K12 strain (Heinrichs *et al.*, 1998), should occupy an area of $R_g \sim 0.4 - 1.0$ nm, and can be represented by $\beta_{\mathrm{OS}} = \Delta\rho V \sim 13 \times 10^{-4}$ nm in case of X-rays ($\Delta\rho$ is the contrast between the SLD of the OS and buffer, and $V$ is the volume of one OS chain). For neutron scattering $\beta_{\mathrm{OS}}$ values should be around $5.6 \times 10^{-4}$ nm, or $-5.1 \times 10^{-4}$ nm in the case of 100 wt% $D_2O$ PBS and full interfacial H/D exchange. Finally, in a first approximation, the scattering contribution of the O-antigen chains can be neglected.

**Periplasm and Peptidoglycan** The periplasmic space – the interstitial layer between the outer and inner membranes – has a thickness of about $20 - 30$ nm and it is less dense than the cytoplasm (Matias *et al.*, 2003). The peptidoglycan, or murein, layer, located within the periplasm is composed of polysaccharides, linear strands of alternating pyranoside-N-acetylglucosamine, and N-acetylmuramic acid. These strands are cross-linked via short sequences of 3-4 amino acids, whose composition is rather well-conserved among strains (Burge *et al.*, 1977). The structure of the peptidoglycan results in a very stiff and rigid network (Pink *et al.*, 2000), having the glycan strands sparsely distributed parallel to the membrane plane (Gan *et al.*, 2008). In *E. coli*, in the exponential growth phase, three of such connected layers with a total thickness of up to than 7.5 nm were reported (Labischinski *et al.*, 1991). Based on its composition and the high hydration of this layer (up to about 90%), we estimate $\rho_{PG}^{\text{X-ray}} \sim 10 \times 10^{-4}$ nm$^{-2}$ and $\rho_{PG}^{\text{neutron}} \sim -0.17 \times 10^{-4}$ nm$^{-2}$, or $5.9 \times 10^{-4}$ nm$^{-2}$ in the case of 100 wt% $D_2O$ suspension.

**Membrane Proteins** In *E. coli*, proteins in the periplasm reach 11 wt% of the dry weight of the totality of bacterial proteins and RNA, and the amount within the cell wall – the membrane protein content – is about 7 wt% (Zimmerman & Trach, 1991). About 50-60 wt% of the cytoplasmic membrane dry mass are membrane proteins, while the outer membrane contains only 5-20 wt% proteins (Seltmann & Holst, 2002). Sizes of these proteins are comparable to those in the cytoplasm, enabling similar approximations for $R_g$ estimates. A small percentage of these proteins is significantly larger, forming complexes that connect either the peptidoglycan layer to the cytoplasmic or outer membrane, or span the full thickness of the cell envelope to form e.g. rotational motors for flagella.

#### *1.3. External cell components*

Finally, *E. coli* possesses a few more components extending outside the cell body. Those include pili, flagella, and the capsule. Pili and fimbriae are hair-like, protein-based, long strands that covers the surface of the cell. They are primarily used for bacterial conjugation and adhesion on surfaces, and they can have a length up to a few tens of nanometers. Flagella enable motility and are also fully composed of proteins, but are, compared to pili and fimbrae, much longer (2 to 12 $\mu$m) hollow tubes with an external diameter of 22 nm (Yamashita *et al.*, 1998; Turner *et al.*, 2012)).

Finally, under certain conditions some cells are able to produce the so-called capsule, which is a thick external layer consisting of a polysaccharide network (Whitfield & Roberts, 1999; Stukalov *et al.*, 2008; Seltmann & Holst, 2002). The chemical composition of the capsule is comparable to the O-antigen of smooth-LPS. Further, the capsule anchoring, is, even in strains having rough-LPS, made up of polysaccharide chains bound to the LPS layer. Detailed structural information of this layer is scarce, however. As such and to the best of our knowledge, no reports on the presence or the type of capsule are available for *E. coli* ATCC 25922. In contrast, K12 strains are known to produce the capsule layer under stress conditions (Seltmann & Holst, 2002).

### 2. Supplementary Figures

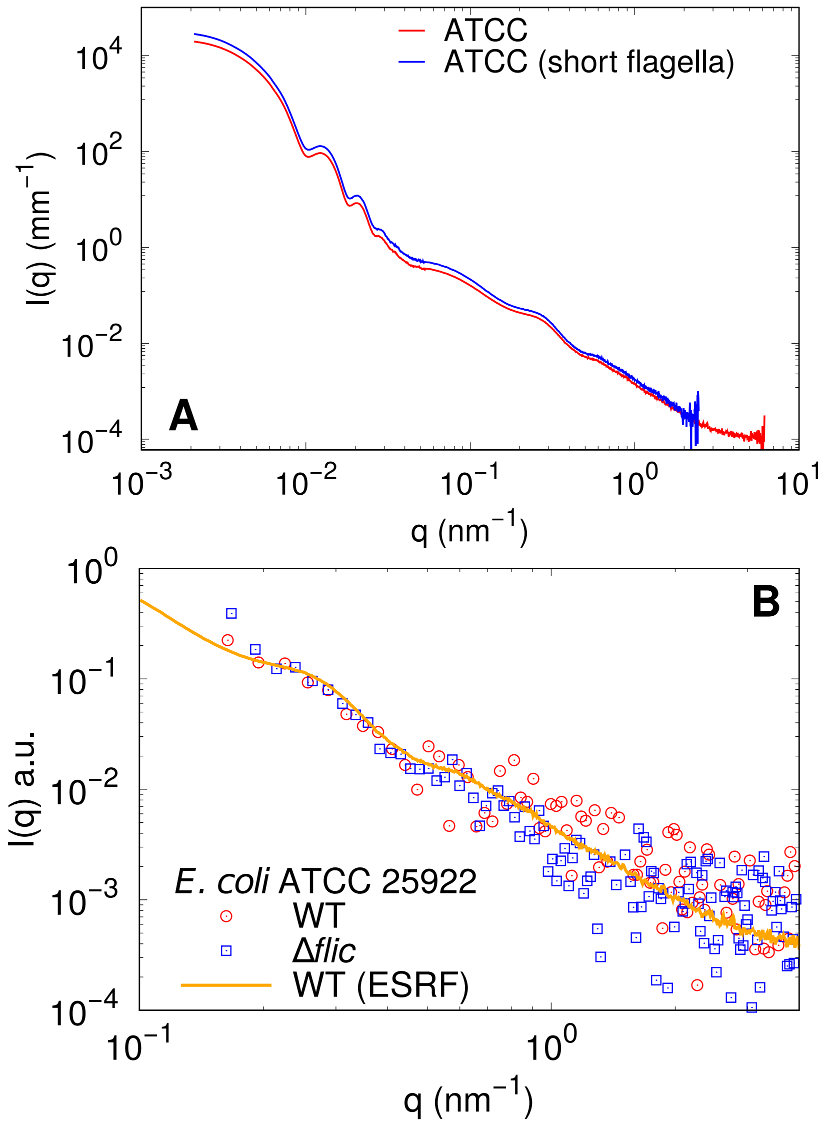


Fig. S1. A) Comparison between scattering intensity curves from *E. coli* ATCC 25922 wild type (WT) (red curve) and from the same cells after fracturing and removing flagella (blue curve). See the main text for details about the sample preparation. B) Comparison between scattering intensity curves from WT (red dots: data acquired in the inhouse SAXS camera, orange line: data acquired at the European synchrotron, ESRF) and from ATCC Δ*fliC* mutant, which does not express flagellin and then does not possess flagella.

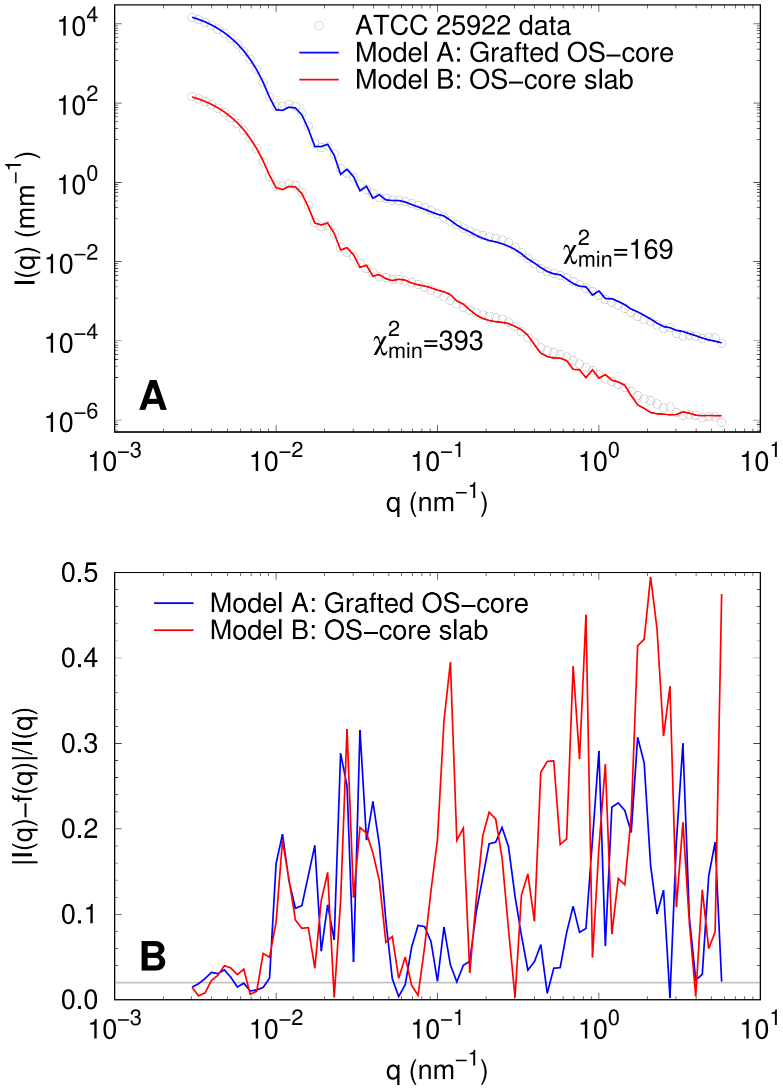


Fig. S2. A) Comparisong of the best fits of USAXS/SAXS data of *E. coli* ATCC 25922 using either grafted polymers to describe the OS cores (model A), or considering the OS-cores via an additional shell, i.e. $I(q) \propto |A_{CS}(q)|^2$ (model B). The two $\chi^2$ values are the minimum values extracted from several Monte Carlo optimization runs. Panel B highlights the corresponding matches of the models in different $q$-ranges. Model B fails in the high-$q$ region ($> 0.1$ nm$^{-1}$).

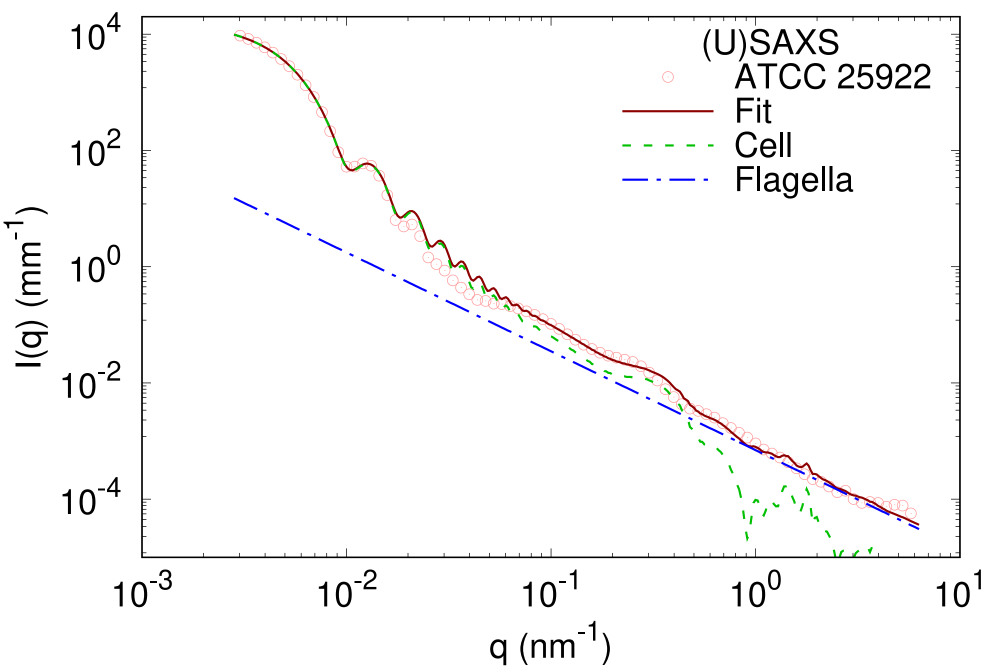


Fig. S3. Example of scattering intensity fitted with the previous model (Semeraro *et al.*, 2017). The power-law describing flagella is able to account for neither the change in effective slope at around $q \sim 0.04$ nm$^{-1}$ nor the scattering feature at about $q \sim 0.1$ nm$^{-1}$.

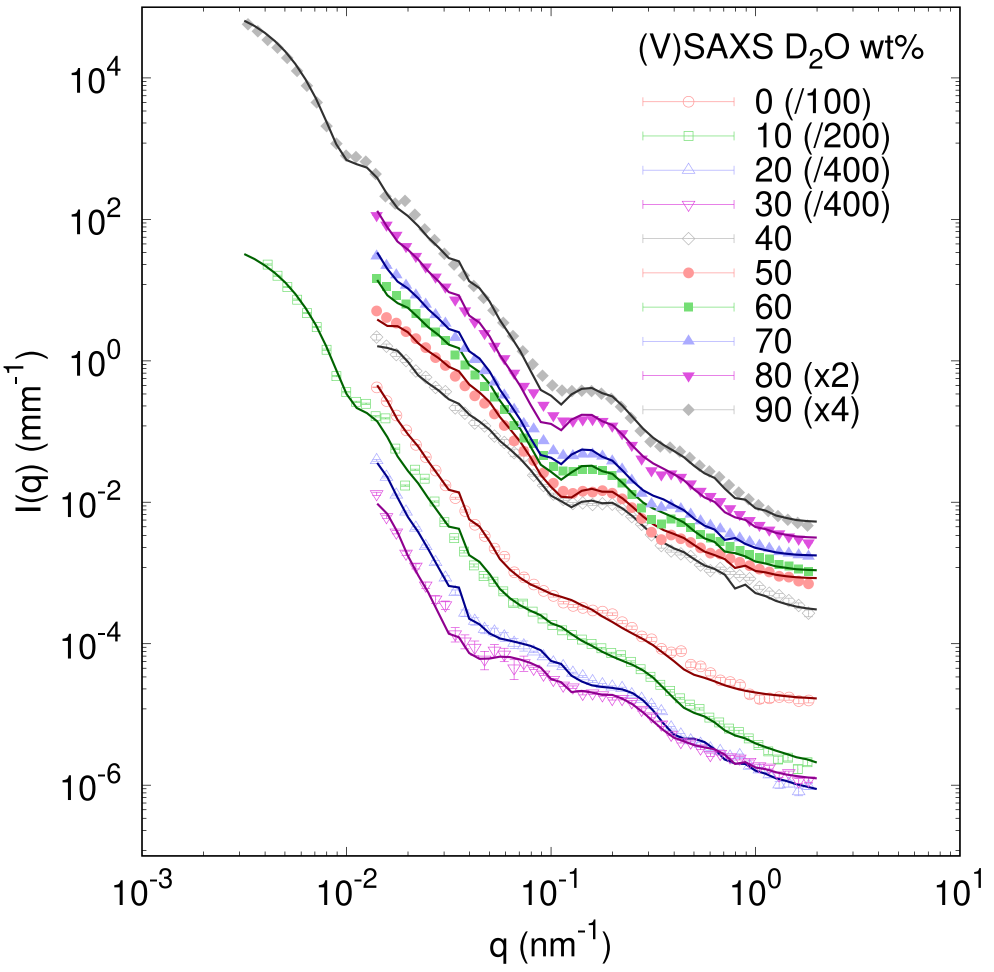


Fig. S4. Contrast-variation VSANS/SANS data in absolute scale and best fits for the strain ATCC 25922 at each $D_2O$ wt%. Scattering curves were scaled for a better visibility.

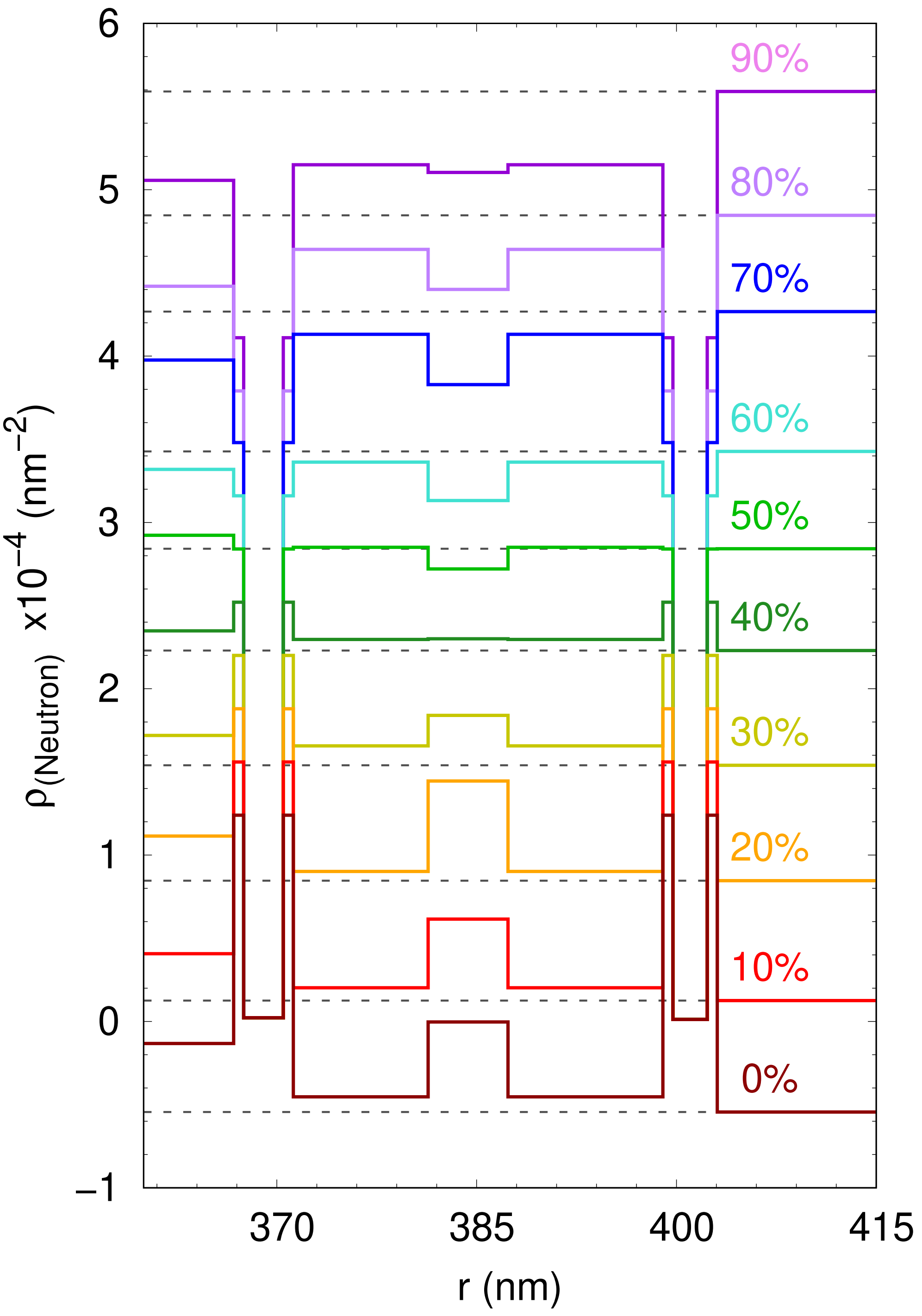


Fig. S5. Neutron SLD profiles of the bacterial ultrastructure of ATCC 25922 strain at different $D_2O$ wt%. It highlights the average positions of both cytoplasmic and outer membrane, and the peptidoglycan layer. *r* describes the distance from the center of the cell along the minor ellipsoidal radius *R*.

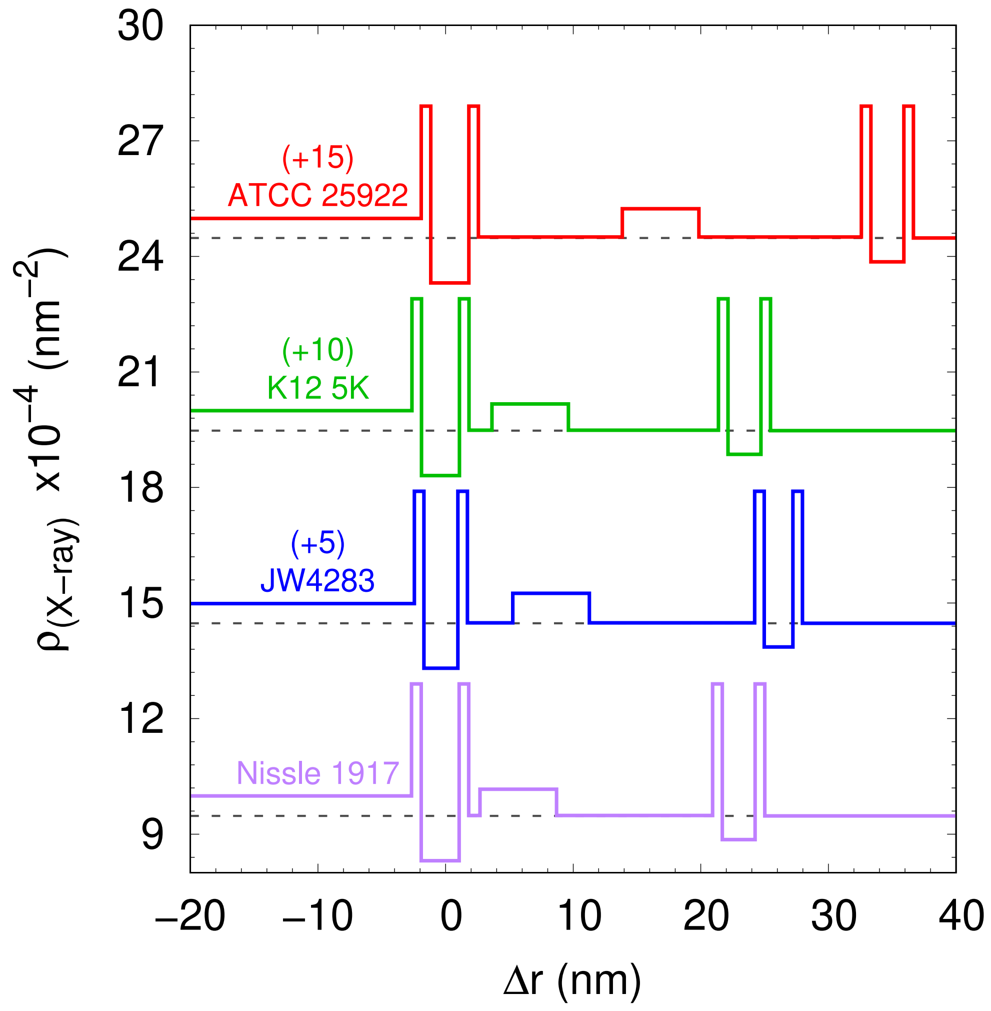


Fig. S6. X-ray SLD profiles of the bacterial ultrastructure of the strains ATCC 25922 (red), K12 5K (green), JW4283 (blue) and Nissle 1917 (purple). $\Delta r$ is the relative distance from the center of mass of the cytoplasmic membrane. SLD profiles were scaled for a better visibility.

### 3. Supplementary Tables

Table S1. *Fixed and fitted local parameters of E. coli ATCC 25922.*

| $\times 10^{-4}$ | USAXS/SAXS / | VSANS/SANS ($D_2O$ wt%) 0 | 10 | 20 | 30 | 40 |
|---|---|---|---|---|---|---|
| $\rho_{CP}$ ($nm^{-2}$) | 9.986±0.011 | -0.132±0.013 | 0.407±0.002 | 1.115±0.010 | 1.719±0.009 | 2.348±0.008 |
| $\rho_{PP}$ ($nm^{-2}$) | 9.503±0.017 | -0.45±0.02 | 0.203±0.012 | 0.902±0.014 | 1.66±0.03 | 2.296±0.017 |
| $\rho_{PG}$ ($nm^{-2}$) | 10.24±0.05 | 0.00±0.04 | 0.62±0.04 | 1.445±0.012 | 1.84±0.06 | 2.30±0.04 |
| $\beta_{OS}$ (nm) | 10.7±0.5 | 4.60±0.18 | 3.79±0.04 | 3.51±0.10 | 2.3±0.2 | 1.67±0.02 |
| $\rho_{ME}$ ($nm^{-2}$)$^a$ | 12.9 | 1.24 | 1.56 | 1.88 | 2.20 | 2.52 |
| $\rho_{TI}$ ($nm^{-2}$)$^a$ | 0.831 | 0.022 | | | | |
| $\rho_{TO}$ ($nm^{-2}$)$^a$ | 0.886 | 0.012 | | | | |
| $\rho_{BF}$ ($nm^{-2}$)$^b$ | 9.476 | -0.544 | 0.1257±0.0007 | 0.846±0.003 | 1.54 | 2.23 |

| $\times 10^{-4}$ | VSANS/SANS ($D_2O$ wt%) 50 | 60 | 70 | 80 | 90 |
|---|---|---|---|---|---|
| $\rho_{CP}$ ($nm^{-2}$) | 2.92±0.05 | 3.32±0.09 | 3.98±0.03 | 4.42±0.03 | 5.06±0.06 |
| $\rho_{PP}$ ($nm^{-2}$) | 2.85±0.05 | 3.362±0.06 | 4.13±0.04 | 4.64±0.04 | 5.15±0.09 |
| $\rho_{PG}$ ($nm^{-2}$) | 2.72±0.04 | 3.13±0.06 | 3.82±0.06 | 4.40±0.06 | 5.10±0.07 |
| $\beta_{OS}$ (nm) | 0.67±0.03 | -0.0019±0.0002 | -0.72±0.09 | -1.73±0.17 | -2.6±0.03 |
| $\rho_{ME}$ ($nm^{-2}$)$^a$ | 2.84 | 3.16 | 3.48 | 3.79 | 4.11 |
| $\rho_{TI}$ ($nm^{-2}$)$^a$ | 0.022 | | | | |
| $\rho_{TO}$ ($nm^{-2}$)$^a$ | 0.012 | | | | |
| $\rho_{BF}$ ($nm^{-2}$)$^b$ | 2.84±0.05 | 3.43±0.06 | 4.27±0.03 | 4.84±0.03 | 5.59±0.06 |

($a$) Fixed values. ($b$) Where the error is noted, the buffer SLD was fitted to account for possible uncertainties in the heavy water percentage.

Table S2. *Fitted contrast values for USAXS/SAXS analysis of ATCC 25922, K12 5K, JW4283 and Nissle 1917 strains.*

| $\times 10^{-4}$ | ATCC 25922 | K12 5K | JW4283 | Nissle 1917 |
|---|---|---|---|---|
| $\rho_{CP}$ ($nm^{-2}$) | 9.986±0.011 | 9.995±0.005 | 9.986±0.009 | 9.993±0.007 |
| $\rho_{PP}$ ($nm^{-2}$) | 9.503±0.017 | 9.486±0.006 | 9.487±0.006 | 9.485±0.004 |
| $\rho_{PG}$ ($nm^{-2}$) | 10.24±0.05 | 10.17±0.02 | 10.25±0.04 | 10.17±0.03 |
| $\beta_{OS}$ (nm) | 10.7±0.5 | 9.9±0.2 | 11.1±0.8 | 10.4±0.4 |
| $\rho_{BF}$ ($nm^{-2}$)$^a$ | 9.476 | | | |

($a$) Fixed parameter.
